# Human MX1 orchestrates the cytoplasmic sequestration of neo-synthesized influenza A virus vRNPs

**DOI:** 10.1101/2024.02.22.581565

**Authors:** Joe McKellar, Francisco García de Gracia, Corentin Aubé, Ana Luiza Chaves Valadão, Marine Tauziet, Mary Arnaud-Arnould, Antoine Rebendenne, Aymeric Neyret, Emmanuel Labaronne, Emiliano Ricci, Bénédicte Delaval, Raphaël Gaudin, Nadia Naffakh, Sarah Gallois-Montbrun, Olivier Moncorgé, Caroline Goujon

## Abstract

Interferon-inducible Myxovirus resistance 1 (MX1) proteins are well-known to restrict influenza A virus (IAV) at early stages during viral replication, impairing the viral transcription/replication process. Herein, we show that this early restriction was only partial against human IAVs, whereas a strong inhibition of viral production was observed. Indeed, relatively high levels of IAV mRNAs and proteins were observed in the presence of human (Hs) and mouse (Mm) MX1 proteins but additional inhibition processes occurred at later stages of IAV life cycle. Hence, MmMx1 induced an abnormal nuclear accumulation of the viral nucleoprotein (NP) at late time points post-infection. This block was also observed, albeit to a much lower extent, with HsMX1. In most HsMX1-expressing cells, vRNPs could be exported from the nucleus to the cytoplasm however a potent defect in subsequent vRNP cytoplasmic trafficking was observed. Indeed, vRNPs were found sequestrated together with cellular co-factors YBX1 and Rab11a in large clusters in the vicinity of the microtubule organization center (MTOC). Live imaging experiments revealed that the transient association of HsMX1 with Rab11a-associated vRNPs favoured their dynein-dependant retrograde transport along microtubules towards the MTOC. Importantly, dynein inhibition prevented the vRNP sequestration and significantly rescued infectious viral production in the presence of HsMX1, showing a significant contribution of these abnormal vRNP clusters in HsMX1 antiviral activity. In conclusion, this study provides the first evidence of IAV vRNPs being re-routed and accumulated away from the plasma membrane, through the coordinated action of HsMX1 restriction factor, dynein and the microtubule network.

## Introduction

Influenza A virus (IAV) is a negative stranded, segmented RNA virus, which is responsible for yearly epidemics of the flu in humans. The IAV genome is composed of eight RNA segments. Each of these viral RNAs (vRNAs) forms a viral ribonucleoprotein complex (vRNP) together with multiple copies of Nucleoprotein (NP) and an RNA-dependent RNA polymerase (RdRp) complex composed of Polymerase Basic 1 and 2 (PB1 and PB2) and Polymerase Acidic (PA) subunits. After internalization of IAV particles through endocytosis and fusion of the viral and endosomal membranes, vRNPs traffic into the nucleus where transcription and replication occur (reviewed in (Simpson & Yamauchi, 2020) and (Hutchinson & Fodor, 2012)). There, viral messenger RNAs (mRNAs) are produced from the vRNA templates by the viral polymerase, a process termed primary transcription. Of note, the M and NS segments give rise to both unspliced and spliced mRNAs (reviewed in (Esparza *et al*, 2022)). Viral mRNAs are then exported to the cytoplasm and translated. Neo-synthesized polymerase subunits and NP are imported to the nucleus to initiate viral genome replication by synthesizing complementary RNA (cRNA) intermediate templates to produce new vRNA molecules. Neo-synthesized polymerase complexes are also responsible for the transcription of more viral mRNAs, a process named secondary transcription (reviewed in (Peacock *et al*, 2019; Fodor & te Velthuis, 2020)). Newly formed vRNPs associate in the nucleus with viral proteins Matrix 1 (M1) and Nuclear Export Protein (NEP) and are exported to the cytoplasm by the cellular Chromosomal Maintenance 1 Homolog (CRM1 / Exportin 1) nuclear export machinery (reviewed in (Lakdawala *et al*, 2016)). vRNPs are subsequently trafficked to an area close to the Microtubule Organization Center (MTOC) (Momose *et al*, 2007; Jo *et al*, 2010; Amorim *et al*, 2011; Kawaguchi *et al*, 2015; de Castro Martin *et al*, 2017). This process is controlled by cellular factors HIV-1 Rev binding protein (HRB) and Y-Box Binding Protein 1 (YBX1) (Eisfeld *et al*, 2011b; Kawaguchi *et al*, 2012). Following this, the PB2 subunit of the vRNP interacts with GTP-bound Rab11a (Veler *et al*, 2022), which mediates vRNP traffic towards the cell surface. Several Rab11-dependant pathways have been proposed to traffic vRNPs towards the plasma membrane, either in a microtubule-dependent or a microtubule-independent manner (Jo *et al*, 2010; Bruce *et al*, 2010; Amorim *et al*, 2011; Eisfeld *et al*, 2011a; de Castro Martin *et al*, 2017; Alenquer *et al*, 2019). Liquid-like phase-separated cytoplasmic inclusions were proposed to be involved in the latter (reviewed in (Amorim, 2019)). Moreover, the endoplasmic reticulum has been shown to be subverted and remodeled by IAV to promote the transport of vRNPs through Rab11-positive irregularly coated vesicles (Castro Martin *et al*, 2017). The vRNPs are then transferred to the plasma membrane by an unelucidated mechanism, where novel viral particles assemble, bud and egress from the host cell.

During IAV infection, interferons (IFN) are produced by infected cells upon viral RNA sensing, leading to the establishment of a so-called antiviral state through the regulation of hundreds of IFN-stimulated genes (ISGs). Several ISGs have been shown to strongly impair IAV replication (reviewed in (McKellar *et al*, 2021)), including the Myxovirus resistance protein 1 (MX1), a member of the Dynamin superfamily of large GTPases (reviewed in (Haller *et al*, 2018)). The protein coded by mouse *Mx1* (MmMx1), the first MX1 gene cloned, is nuclear and probably the most potent restriction factor against IAV identified to date (Krug *et al*, 1985; Staeheli *et al*, 1986; Pavlovic *et al*, 1992). Human MX1 (HsMX1), which is ∼65% identical to MmMx1 at the aminoacid level, is solely cytoplasmic and inhibits viruses replicating in the cytoplasm such as LaCrosse virus (LACV), as well as viruses replicating in the nucleus, like IAV (Frese *et al*, 1996; Kochs *et al*, 2002). Remarkably, HsMX1 is actually able to broadly inhibit the replication of a vast range of RNA and DNA viruses (reviewed in (Verhelst *et al*, 2013)). In terms of IAV inhibition, MmMx1 and HsMX1 were shown to inhibit different stages of the replication cycle (Pavlovic *et al*, 1992). However, it is generally accepted that both proteins act on IAV transcription/replication. MmMx1 targets primary transcription, potentially by affecting elongation of viral transcripts and/or by interfering with the interaction between NP and PB2 (Krug *et al*, 1985; Pavlovic *et al*, 1992; Verhelst *et al*, 2012). HsMX1 inhibits viral replication after primary transcription, limiting viral genome amplification (Pavlovic, Haller, et Staeheli 1992; Nigg et Pavlovic 2015). Interestingly, longer IAV segments are more strongly inhibited by MX1 proteins than their shorter counterparts (Krug *et al*, 1985; Pavlovic *et al*, 1992). NP was shown to be an important determinant for MX1 sensitivity as avian IAVs are more sensitive to HsMX1 restriction than human IAVs, and this was mapped to NP (Dittmann *et al*, 2008; Zimmermann *et al*, 2011; Mänz *et al*, 2013). However, the detailed molecular mechanisms through which MX1 proteins exert their anti-IAV activity remain to be fully elucidated. Interestingly, HsMX1 inhibits LACV replication without impacting initial transcription of the viral genome or Nucleoprotein (LACV-N) expression (Frese *et al*, 1996). Instead, HsMX1 specifically binds to, and aggregates LACV-N into large, perinuclear complexes (Kochs *et al*, 2002). As a consequence of this sequestration, LACV genome amplification is severely inhibited. Such nucleoprotein aggregation phenotypes were also observed for other bunyaviruses (Kochs *et al*, 2002; Andersson *et al*, 2004). Furthermore, HsMX1 was shown to be recruited to replication factories of African swine fever virus (ASFV), a large DNA virus, in the perinuclear region, where it surrounded them (Netherton *et al*, 2009). In the case of IAV, such a relocalization of MX1 proteins and colocalization with viral proteins have never been reported but late time points post-infection were usually not considered, as the known restrictions intervene early in the viral life cycle.

In the present study, we investigated whether MX1 proteins could affect later stages of IAV replication cycle in addition to the inhibitions exerted on viral transcription/replication. Using human IAVs, we confirmed that MmMx1 and HsMX1 proteins did indeed reduce IAV transcription/replication. However, this inhibition was only partial, with a surprisingly strong expression of some viral mRNAs and proteins (including NP), at late stages of infection. Interestingly, we showed that in the vast majority of MmMx1-expressing cells, NP was expressed but localized abnormally to the nucleus at a late time point post-infection, when it should be cytoplasmic. In contrast, in most HsMX1-expressing cells, NP could be exported to the cytoplasm under the form of vRNPs, but further cytoplasmic trafficking was prevented and vRNPs were abnormally accumulated together with Rab11a-positive vesicles in the vicinity of the MTOC. This inhibition of vRNP trafficking could potentially be directly mediated by HsMX1, as the antiviral protein was transiently found associated with the abnormal vRNP clusters. Microtubule inhibition with nocodazole led to small and dispersed vRNP clusters throughout the cytoplasm, with which HsMX1 remained associated overtime. In contrast, the dynein inhibitor Dynapyrazole-A limited the formation of the abnormal vRNP clusters and significantly rescued infectious viral production in the presence of HsMX1. Altogether, these data show that HsMX1 prevents the trafficking of IAV vRNPs towards the plasma membrane by inducing their dynein-dependant retrograde transport and their sequestration in the vicinity of the MTOC. To our knowledge, these data provide the first evidence of HsMX1 directly perturbing the late stages of IAV replication.

## Materials and methods

### Plasmid constructs

The pRRL.sin.cPPT.SFFV/IRES-puro.WPRE lentiviral vector system has been described previously (Doyle *et al*, 2018), as have pRRL.sin.cPPT.SFFV/IRES-puro.WPRE-FLAG-Renilla, -FLAG-HsMX1 and FLAG-MmMx1 (McKellar *et al*, 2023). The eGFP coding sequence (CDS) along with a multiple cloning site (MCS) was cloned using NotI/XhoI restriction sites to generate pRRL.sin.cPPT.SFFV/eGFP:MCS.IRES-puro.WPRE. Rab11a and Rab11a-S25N CDS were PCR-amplified from plasmids obtained from Dr Cécile Gauthier-Rouvière (CRBM, Montpellier), and cloned into the pRRL.sin.cPPT.SFFV/eGFP:MCS.IRES-puro.WPRE backbone to obtain pRRL.sin.cPPT.SFFV/eGFP:Rab11a.IRES-puro.WPRE and pRRL.sin.cPPT.SFFV/eGFP:Rab11a-S25N.IRES-puro.WPRE using BamHI and XhoI sites. Rab11a-Q70L point mutant was generated by site-directed mutagenesis and inserted into pRRL.sin.cPPT.SFFV/eGFP:MCS.IRES-puro.WPRE. H2B-FAST was a gift from Arnaud Gauthier (Addgene: #130722) (Plamont *et al*, 2016). FAST CDS was flanked with an N-terminal FLAG-tag sequence and a C-terminal multiple cloning sequence (MCS), and cloned into pRRL.sin.cPPT.SFFV/IRES-puro.WPRE to produce pRRL.sin.cPPT.SFFV/FLAGFAST-MCS-IRES-puro.WPRE. HsMX1 coding sequence was sub-cloned into both pRRL.sin.cPPT.SFFV/eGFP-MCS-IRES-puro.WPRE and pRRL.sin.cPPT.SFFV/FLAGFAST-MCS-IRES-puro.WPRE backbones. Constructs of interest were also cloned into pRRL.sin.cPPT.SFFV/IRES-blasti.WPRE and pRRL.sin.cPPT.SFFV/IRES-neo.WPRE.

IAV H3N2 A/Victoria/3/75 Pol I reverse genetic plasmid pPolI-RT-Vic-PA-FLAG was obtained by introducing two FLAG-tag sequences in 3’ of the PA coding sequence following the strategy we used previously (Doyle *et al*, 2018). The reverse genetics plasmids for influenza virus A/WSN/33 (Fodor *et al*, 1999) were initially kindly provided to S. Munier and N. Naffakh by G. Brownlee (Sir William Dunn School of Pathology, Oxford, UK). The pPolI-PA-LL-mScarlet plasmid was obtained by substituting the LL-Gluc1 sequence lying between the NotI and SpeI sites in the pPolI-PA-LL-Gluc1 plasmid, as done previously for other reporter systems (Munier *et al*, 2013).

For transient transfection experiments, previously described pCAGGS expression plasmids were used (McKellar *et al*, 2023).

All plasmid constructs were verified by Sanger sequencing (Eurofins) or Oxford Nanopore sequencing (Plasmidsaurus).

### Cell lines

Cell lines (A549 (human lung adenocarcinoma; ATCC CCL-185), Human Embryonic Kidney 293T (HEK293T; ATCC CRL-3216) and Madin-Darby canine kidney (MDCK; ATCC CCL-34)) were grown in Dulbecco’s modified Eagle medium (DMEM) supplemented with 10% fetal bovine serum (FBS), 100 μg/mL penicillin and 100 units/mL streptomycin.

### Influenza virus production and infection

A/Victoria/3/75 (H3N2), A/Victoria/3/75-PA-FLAG (A/Victoria/3/75 modified to code for PA tagged with two FLAG tags in C-terminus), A/Victoria/3/75-NLuc (IAV-NLuc, A/Victoria/3/75 containing the Nanoluciferase (NLuc) coding sequence in the PA segment) and A/WSN/33 (H1N1) were produced as described previously (Goujon *et al*, 2013). A/WSN/33 reverse genetic system (Fodor *et al*, 1999) was a kind gift from Prof. Sandie Munier (Institut Pasteur, Paris). A/WSN/33-PA:mScarlet was produced in Dr Nadia Naffakh’s lab as described previously for other reporter systems (Munier *et al*, 2013). Viral stocks were titrated by plaque assays on MDCK cells.

For the growth curve experiments, A549 cells stably expressing FLAG-tagged Renilla, HsMX1 or MmMx1 were seeded in 12 well plates and were infected for 1 h at MOI 0.005 in technical duplicates. Cells were washed with PBS and incubated at 37°C in 1 ml of serum-free DMEM containing 1 µg/ml TPCK-treated trypsin. Samples were collected at 24, 48, 72 and 96 h post-infection for viral titration by plaque assays on MDCK cells. For IAV-NLuc reporter virus assays, cells were infected at MOI 1 for 24 h and cells were frozen dry at -80°C for 30 min and lysed in Passive Lysis Buffer (Promega). Bioluminescence was measured using the Nano-Glo® Luciferase Assay System (Promega) on a Tecan plate reader and results were analysed using GraphPad PRISM.

When indicated, cycloheximide (CHX) treatment was performed by incubating cells with pre-warmed complete media containing CHX (50 µg/mL) following the 1 h incubation with IAV and for the remainder of the infection. Furthermore, for experiments using Nocodazole (Sigma-Aldrich) or Dynapyrazole-A (Dyn-A) (Sigma-Aldrich), media was changed either 3 h or 21 h post-infection and replaced with pre-warmed media containing 50 µM Nocodazole or 40 µM Dyn-A for the remainder of the infection, as indicated.

For the viral rescue experiments using Dyn-A, HsMX1- or control-expressing cells were infected with IAV (MOI 1) in serum-free medium, which was replaced 1 h later with medium containing serum (complete media). 3 h post-infection, the culture medium was replaced again with complete media supplemented with 40 µM Dyn-A or the equivalent volume of DMSO (of note, we observed that Dyn-A precipitates in serum-free medium, therefore the incubation was done in complete media). As Dyn-A inhibition of dynein was shown to be only partially reversible in the absence of serum in the medium (Steinman *et al*, 2017), the medium was replaced again at 24 h post-infection with serum-free medium devoid of DMSO or Dyn-A but containing 1 µg/ml TPCK-treated trypsin (or not, for an immunofluorescence analysis control). The supernatants were harvested at 48 h and 72 h post-infection (i.e., 24 h and 48 h post serum removal) and the infectious viruses were titrated by plaque assays.

### Lentiviral vector production and transduction

Lentiviral vector (LV) stocks were obtained by polyethyleneimine (PEI) or Lipofectamine 3000 (Thermo Scientific)-mediated transfection of HEK293T cells with vectors expressing Gag-Pol (p8.91), VSV-G (pMD.G) (Naldini *et al*, 1996) and the indicated miniviral genome at a ratio of 1:0.25:1. The culture medium was renewed 8 h post-transfection and harvested 2 days later. Cells were transduced with filtered supernatants for 8 h prior to media replacement. After 36 h, the relevant antibiotics (Puromycin, G418, Sigma-aldrich; blasticidin, InVivoGen) were added to select the transduced cells.

### Immunoblotting analysis

For immunoblotting, cells were washed in phosphate buffered saline (PBS) 1X and lysed in deoxycholate (DOC)-containing lysis buffer with 1X Laemmli [10 mM Tris-HCl pH7.6, 150 mM NaCl,1% Triton X100, 1 mM EDTA, 0,1% DOC, 2% SDS, 5% Glycerol, 100 mM DTT, 0,02% bromophenol blue] followed by denaturation. SDS-PAGE was performed and subsequently analysed by immunoblotting. The following primary antibodies (and dilutions) were used: anti-M1 (GeneTex, GTX127356, 1/1000), NS1 (GeneTex, GTX125990, 1/1000), PB2 (GeneTex, GTX125926, 1/1000), NP (Southern Biotech, 10770-01, 1/1000), M2 (GeneTex, GTX125951, 1/1000), H2B (Sigma-Aldrich, 07-317, 1/1000), Actin (Sigma-Aldrich, A1978, 1/5000). Primary antibodies were incubated for 1 h at room temperature (RT) or overnight at 4°C, followed by incubation with HRP-conjugated secondary antibodies (anti-Rabbit-HRP, Life Technologies A24537, 1/2500; anti-Mouse-HRP, Life Technologies, A24512, 1/2500) for 1 h at RT. Some antibodies used were directly coupled to HRP and incubated 1 h at RT (FLAG-HRP, Sigma-Aldrich, A8592, 1/5000; Myc-HRP, Sigma-Aldrich, A5598, 1/5000; GAPDH-HRP, Sigma-Aldrich, G9295, 1/5000). Bioluminescence emission was subsequently measured using Clarity ECL Western Blotting Substrate (Bio-Rad) and a ChemiDoc system (Bio-Rad).

### Nucleocytoplasmic fractionations

Cells were plated the day prior to infection in 6 well plates and grown to sub-confluency, infected or not for 24 h then washed with ice-cold PBS 1X. Cells were harvested, pelleted and resuspended in ice-cold PBS 1X. Cells were pelleted at 10 000 rpm for 7 seconds and resuspended in 900 µl of ice-cold fractionation buffer [PBS 1X, 0.1% IGEPAL CA-630 (Euromedex), cocktail of protease inhibitors (Roche)] and pipetted up and down several times. An aliquot corresponding to the whole cell lysate (WCL) was harvested (300 µL), and 100 µL of Laemmli-DOC 4X was added and the WCL stored on ice. The remaining volume of lysate was cleared at 10 000 rpm for 7 seconds and 300 µL of supernatant was taken, corresponding to the cytoplasmic fraction, and supplemented with 100 µL of Laemmli-DOC 4X. The remaining supernatant was discarded and the pellet washed once with ice-cold fractionation buffer and pelleted at 10 000 rpm for 7 seconds, the supernatant discarded. The pellet was resuspended in 800 µL of Laemmli-DOC 1X, corresponding to the nuclear fraction. All fractions were boiled for 10 min and resolved by immunoblotting as described above.

### Immunofluorescence, FISH and confocal microscopy

Cells stably expressing the constructs of interest were plated on coverslips in 24-well plates and grown to sub-confluency. For HEK293T cells, coverslips were pre-coated with Poly-L-Lysine (Sigma-Aldrich). Cells were infected or not with the indicated viruses and fixed with either 2% paraformaldehyde (PFA) in PBS 1X for 10 min at RT or ice-cold Methanol at -20°C for 20 min. When PFA was used, permeabilization was performed by incubation in 0.2% Triton X-100 in PBS 1X for 10 min at RT. Quenching and blocking in NGB buffer [50 mM NH4Cl, 2% Goat serum, 2% Bovine serum albumin, in PBS 1X] for 1 h followed. The following primary antibodies were used: anti-FLAG (Sigma-Aldrich, F1804, 1/1000), Myc (ProteinTech, 16286-1-AP, 1/150), M1 (GeneTex, GTX127356, 1/100), NS1 (GeneTex, GTX125990, 1/250), PB2 (GeneTex, GTX125926, 1/500), NP (either from Southern Biotech, 10770-01, 1/100, or supernatant from hybridoma HB-65, H16-L10-4R5, ATCC, 1/100), M2 (GeneTex, GTX125951, 1/100), NS2/NEP (GeneTex, GTX125953, 1/150), TGN46 (BioRad, AHP500, 1/100), YBX1 (ProteinTech, 20339-1-AP, 1/50), Rab11 (Cell Signaling, 5589 1/50), Rab11a (Proteintech, 20229-1-AP, 1/50), α-Tubulin-FITC (Sigma-Aldrich, F2168, 1/300), HRB (ProteinTech, 12670-1-AP 1/100), and Pericentrin (Abcam, ab4448, 1/100). Primary antibodies were incubated for 1 h followed by washes and incubation with Alexa Fluor conjugated secondary antibodies (Thermofisher Scientific) for 1 h. When needed, primary antibodies were conjugated to Alexa Fluor with the Zenon labelling kits (Thermofisher Scientific). Cells were then incubated with DAPI (ThermoFisher Scientific) for 15 min or Filipin (100 µg/mL, Sigma-Aldrich, F4767, a kind gift from Dr María Moriel Carretero, CRBM) and mounted on glass slides using ProLong™ Gold Antifade Mountant (Thermofisher Scientific).

For FISH experiments, A/Victoria/3/75 M1 vRNA probes were designed (27 total; biosearchtech.com/stellaris-designer) and amine-modified (5AmMC6) oligonucleotides were synthesized (Eurofins). Oligonucleotides were resuspended, chloroform extracted thrice to remove any free amines, ethanol precipitated and resuspended at 25 µg/µL. 50 µg of the oligonucleotide pool was labelled using the Alexa Fluor 488 Oligonucleotide Amine Labeling Kit (ThermoFisher Scientific) as per the kit instructions. Labelled oligonucleotides were cleaned and concentrated using the Oligo Clean & Concentrator kit (Zymo Research). Finally, probes were adjusted to a final concentration of 12.5 µM and stored at -20°C. For FISH coupled to immunofluorescence experiments, cells were plated and grown to sub-confluency on coverslips in 24-well plates. Cells were infected (or not) for 24 h with A/Victoria/3/75 at MOI 1 and fixed for 10 min in 2% PFA. Cells were permeabilized 1 h in 70% EtOH and washed with Stellaris RNA FISH Wash Buffer A (BioSearch). Cells were incubated cell side down in 100 µL of Stellaris RNA FISH Hybridization Buffer (BioSearch) containing M1 vRNA probes (1/100) and the primary antibodies of interest (NP, Southern Biotech, 10770-01, 1/100) and incubated in a humid chamber at 37°C in the dark for 16 h. Following washes, incubation was performed with Alexa Fluor conjugated secondary antibodies in Stellaris RNA FISH Wash Buffer A for 30 min in a humid chamber at 37°C in the dark and subsequent DAPI incubation in the same conditions was performed. A final wash with Stellaris RNA FISH Wash Buffer B (BioSearch) was performed before mounting and imaging as above.

Imaging was performed on either an LSM880 confocal microscope with an Airyscan module (Zeiss) or an LSM980 8Y with Airyscan 2 module (Zeiss), using a 63x lens. Pre-processing of the raw Airyscan images was performed on the ZEN Black or ZEN Blue softwares. Post-processing of all images was performed using the FIJI software (Schindelin *et al*, 2012).

### Live cell imaging

Cells stably expressing the constructs of interest were plated in Cellview glass bottom dishes (Greiner), or 384-well plateq with glass bottoms (Dutscher), grown to sub-confluency and infected or not. Media was changed to DMEM without phenol red (ThermoFisher Scientific), supplemented with L-Glutamine, Pyruvate, 10% FBS, 100 μg/mL penicillin, 100 units/mL streptomycin and, when specified, with nocodazole (50 µM). For FAST:HsMX1 live imaging, N871b (a red-shifted FAST substrate, a kind gift from Dr Mikhail Baranov (IBCH, Moscow, Russia) (Povarova *et al*, 2019)), at 1/250, was added to the media prior to imaging. Cells were imaged at the indicated times post-infection using a Dragonfly spinning disk confocal microscope (Andor) with a 100X lens. Images were post-processed using the FIJI software and representative frames were chosen from time courses for figures.

All movies have been deposited on FigShare:

https://figshare.com/articles/media/Movies_from_McKellar_et_al_bioRxiv_2024_Human_MX1_induces_the_cytoplasmic_sequestration_of_neo-synthesized_influenza_A_virus_vRNPs/25289647

### Image quantification

For the quantifications, the images were acquired on fields of cells that were selected at random on the coverslips. All images were post-processed and minimum and maximum intensity levels of the channels for DAPI, NP and any other studied protein (when applicable) were homogenized (using the Brightness/Contrast Tool of FIJI) to the same value within a given experiment for FLAG-tagged Renilla-, HsMX1- and MmMx1-expressing cells. The number of nuclei per field was assessed and the number of cells containing a positive signal for NP were counted manually. The NP-positive cells were classified into three categories: exclusive nuclear localization, exclusive cytoplasmic localization and hybrid nuclear and cytoplasmic localizations. Further quantification was then performed to determine the percentage of NP-positive cells that contained NP perinuclear accumulations in FLAG-HsMX1-expressing cells. For experiments determining the presence or absence of a given protein (or vRNA) from the NP-positive perinuclear accumulations in FLAG-HsMX1-expressing cells, all channel minimum and maximum intensities were homogenized to the same values within a given experiment (using the Brightness/Contrast Tool of FIJI) and reference images of presence and absence phenotypes were selected and used as benchmarks to quantify the phenotypes accordingly. The number of fields and average number of cells analysed for each experiment are indicated in the figure legends.

### Electron microscopy

A549 cells expressing either FLAG-HsMX1 or FLAG-Renilla were infected or not with A/Victoria/3/75 at MOI 1 for 24 h and fixed with 2.5% (v/v) glutaraldehyde in PHEM buffer [60 mM PIPES; 25 mM HEPES; 10 mM EGTA; 2 mM MgCl_2_] and post-fixed in osmium tetroxide 1% / K_4_Fe(CN)_6_ 0,8%, at RT for 1 h. The samples were then dehydrated in successive ethanol baths (50/70/90/100%) and infiltrated with propylene oxide/EMbed812 mixes before embedding. 70 nm ultrathin cuts were made on a PTXL ultramicrotome (RMC, Tucson, AZ, USA), stained with uranyle acetate/lead citrate and observed on a Tecnai G2 F20 (200 kV, FEG) transmission electron microscope at the Electron Microscopy Facility COMET, INM, Montpellier.

### RNA quantification by RT-qPCR

0,5-2 × 10^6^ cells were collected 8 h or 24 h following mock infection or infection with A/Victoria/3/75 infection at MOI 1 and total RNAs were isolated using the RNeasy kit with on-column DNase treatment (Qiagen). cDNAs were generated using 125 ng RNA (High-Capacity cDNA Reverse Transcription Kit using 250 nM oligodT instead of the provided random primers, ThermoFisher Scientific, 4368813) and analysed by quantitative (q)PCR. The following primers and probes, specific for PA and M1, were used: PA-forward 5’-TTGCTGCACAGGATGCATTA-3’, PA-reverse 5’-AGATTGGAGAAGACGTGGCT-3’, PA-probe 5’-[FAM]-TGGCTCTGCAATGGGACACCTCTGC-[TAMRA]-3’, M1-forward 5’-CAGCACTGGAGCTAGGATGA-3’, M1-reverse 5’-AAGGCTATGGAGCAAATGGC-3’, M1-probe 5’-[FAM]-TGGCCTCTGCTGCCTGCTCA-[TAMRA]-3’, M2-forward 5’-GAGGTCGAAACGCCTATCAG-3’. The corresponding pPolI-RT-Victoria plasmids (a gift from Prof. Wendy Barclay) were used to generate a standard curve and ensure the linearity of the assay (detection limit: 10 molecules per reaction).

### Oxford Nanopore long read sequencing

RNAs from A549 cells expressing FLAG-tagged Renilla, HsMX1 or MmMx1 and infected or not with IAV at MOI 1 were extracted using the miRNeasy Mini kit (Qiagen). RNA quality was analysed using a BioAnalyzer (Agilent) and the Agilent RNA 6000 Nano Kit. RNAs with an RNA integrity number (RIN) higher than 9.0 were used for library preparation. Next, poly-A(+) RNAs were isolated using oligo (dT)25 Dynabeads (Invitrogen) and 200 ng were used for library preparation using the Direct cDNA Native Barcoding kit (SQK-DCS109 with EXP-NBD104) replacing the VN primer by a mix of eight primers, with each primer targeting one of the eight viral mRNAs (see Supplementary Table 1 for primer sequences). For primer design, the poly-T sequence and the last two nucleotides of the VN primer were replaced by a sequence that recognizes specifically the 3’ end of each viral mRNA (indicated in red in Supplementary Table 1). The final concentration of each primer in the reverse transcription mix was 0.05 µM. As a control, an extra primer targeting the 3’end of the GAPDH mRNA was added to the reverse transcription reaction. The obtained library was loaded into a flow cell (Flo-MIND106D) and sequenced using a Minion Mk1B device using the MinKNOW (21.11.7) software. The basecalling of the data was performed using the MinKNOW (21.11.7) software with the High Accuracy option.

For RNA sequencing data analysis, the reads with an extension lower than 15 bp were eliminated and the alignment was performed using Minimap2 (V2.17) (Li, 2018) (specific command line: -ax splice) against a merged version of the *Homo sapiens* genome (GRCh38.p13) with the sequences of the A/Victoria/3/75 mRNAs based on the reverse genetic plasmids. Alignment files were converted into BAM format, sorted and indexed using SAMtools version 1.6 (Li *et al*, 2009). Cellular and viral transcript quantification was realized using IsoQuant version 3.3.0 (Prjibelski *et al*, 2023) with a merge annotation file of the Homo sapiens genome (gencode version 38 basic annotation) and an adapted version from the NCBI of A/Victoria/3/75 (OFL_ISL_105614). One sample t-test was performed with GraphPad Prism.

Of note, due to the very short length of M3 mRNA (291 nt) and the automatic trimming of the 5’ ends of the sequences, unambiguous alignements of reads were particularly difficult. M3 mRNAs were thus not quantified in the analysis.

### Statistical analyses

All statistical analyses were performed with the GraphPad Prism software. The analysis types are indicated in the figure legends. Comparisons are relative to the indicated condition. P-values are indicated as the following: ns = not significant, p<0.05 = *, p<0.01 = **, p<0.001 = *** and p<0.0001 = ****.

## Results

### IAV RNAs and proteins accumulate in infected cells despite the presence of MX1 proteins

In order to explore the stages of IAV life cycle inhibited by MX1 proteins, we generated stable HsMX1-, MmMx1- and control-(Renilla-)expressing A549 cell lines, as described previously (McKellar *et al*, 2023). To assess the antiviral phenotypes imposed by MX1 proteins in these cell lines, we measured infectious IAV production following infection with either A/Victoria/3/75 (H3N2) or A/WSN/33 (H1N1) at MOI 0.005 (Fig. 1A). As expected, HsMX1 inhibited replication of both IAV strains (∼1-2.5 log inhibition as compared to the control), however these strains were able to replicate to some extent in the presence of HsMX1. In contrast, there was very little to no infectious virus produced from MmMx1-expressing cells, thereby confirming the particularly potent anti-IAV activity of MmMx1 (Zimmermann *et al*, 2011). Next, we evaluated viral protein accumulation by immunoblotting, following single-round infection of the different cell lines with A/Victoria/3/75 (Fig. 1B) (or A/Victoria/3/75 bearing a FLAG-tagged PA, Fig. S1A) at MOI 1 (a MOI at which MX1 proteins still exert a profound antiviral effect, Fig. S1B). At 8 h post-infection, PB2, M1 and NS1 viral proteins could be detected in the control cells, while M2 and NP were not detectable. No obvious impact of HsMX1 was observed, whereas the amounts of viral proteins detected were systematically lower in MmMx1-expressing cells. At 24 h post-infection, all the tested viral proteins were detected in the control cells, and a cycloheximide (CHX) control confirmed that they had been neo-synthesised. In the presence of MmMx1, an inhibition of PB2, NP, M2 and PA accumulation was observed, while the impact on M1 and NS1 was minor (Fig. 1B and S1A). Again, at this time point, HsMX1 had very little to no impact on viral protein accumulation (Fig. 1B and S1A). Similar results were obtained with A/WSN/33 infection (Fig. 1C). In line with the western blot data, RT-qPCR analyses confirmed that A/Victoria/3/75 PA mRNA accumulation was more strongly inhibited by MX1 proteins than that of M1 mRNA (Fig. S1C). These results confirmed that, in agreement with the literature (Pavlovic *et al*, 1992), viral mRNAs and proteins issued from the longest segments (PB2, PB1, PA) were more strongly inhibited by MX1 proteins than those from the shortest segments (NS, M).

**Figure 1.**
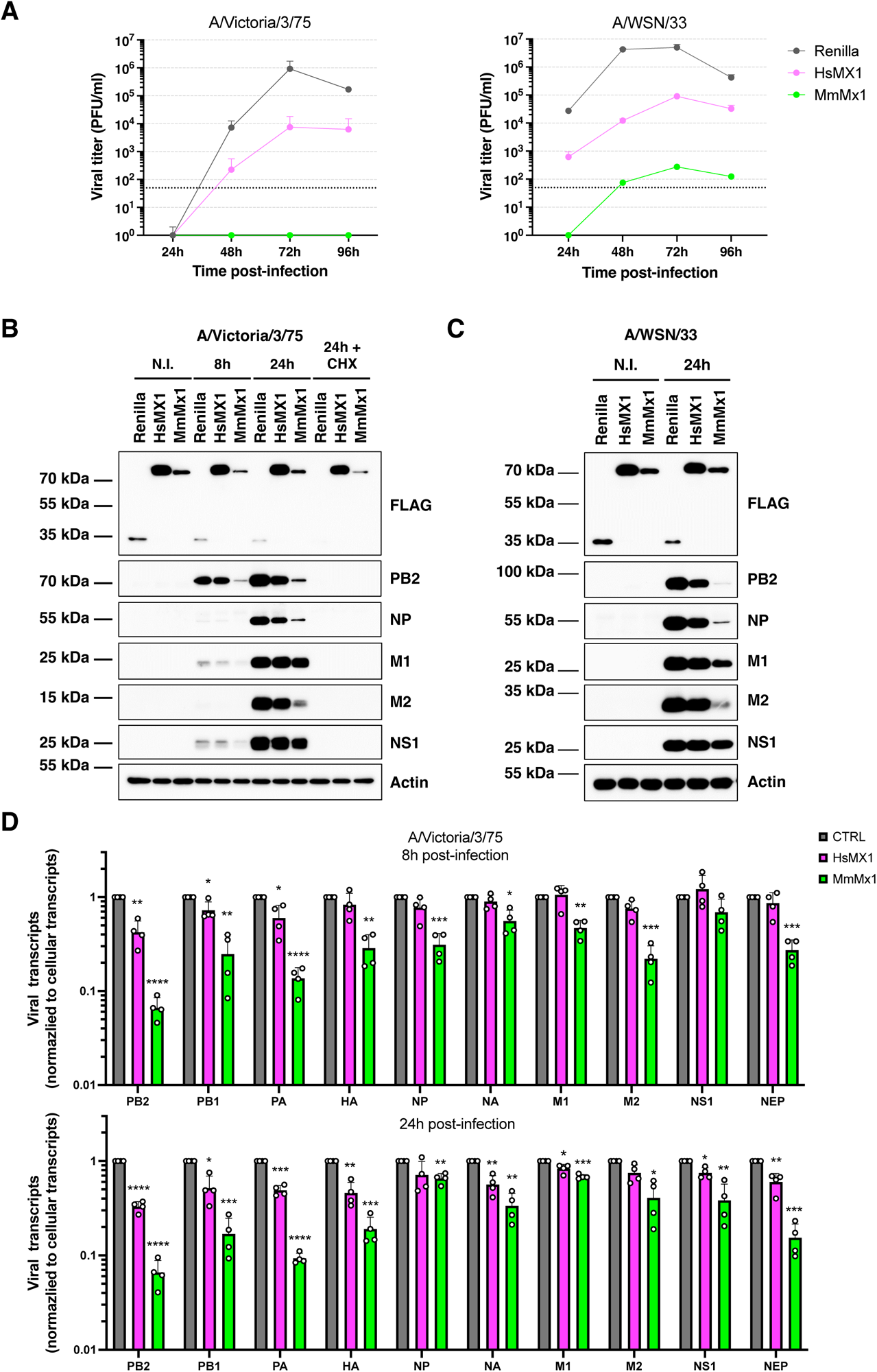
MX1 proteins inhibit early stages of IAV infection and preferentially impact longer segments. **A.** A549 cells expressing FLAG-tagged Renilla luciferase, HsMX1 or MmMx1 were infected with A/Victoria/3/75 (left panel) or A/WSN/33 (right panel) at MOI 0.005. The supernatants were harvested at the indicated time points post-infection and infectious virus production was measured by plaque assays on MDCK cells. The data correspond to a representative experiment, with the means and standard deviations (SD) of technical triplicates. The dotted line represents the limit of detection. PFU: plaque forming units. **B.** A549 cells expressing FLAG-tagged Renilla, HsMX1 or MmMx1 were infected or not (N.I.) with A/Victoria/3/75 at MOI 1, in the presence or not of CHX, lysed at the indicated time points and viral protein accumulation analysed by immunoblot; Actin served as a loading control. Representative immunoblots (out of 3 independent experiments) are shown. CHX: cycloheximide. **C.** A549 cells expressing FLAG-tagged Renilla, HsMX1 or MmMx1 were infected with A/WSN/33 at MOI 1, lysed 24 h later and viral protein accumulation analysed by immunoblot; Actin served as a loading control. Representative immunoblots (out of 2 independent experiments) are shown. **D.** A549 cells expressing FLAG-tagged Renilla (CTRL), HsMX1 or MmMx1 were infected with A/Victoria/3/75 at MOI 1, the RNAs were extracted at 8 h (top) and 24 h (bottom) post-infection and Oxford Nanopore long read sequencing analysis was performed on IAV cDNAs. Data represented are from 4 independent experiments.

Remarkably, the accumulation of M2 protein, which is translated from spliced mRNAs obtained from the short segment M, was more affected by MmMx1 than that of M1, which is translated from unspliced M mRNAs (Fig. 1B, 1C, S1A). To address a potential impact of MX1 proteins on spliced mRNAs and to determine whether IAV protein abundance correlated or not with mRNA abundance, we performed Oxford Nanopore long-read sequencing on IAV reverse transcribed cDNAs, at 8 h and 24 h post A/Victoria/3/75 infection. The amounts of viral transcripts normalized to the cellular transcripts revealed a good correlation with the RT-qPCR and immunoblot data (Fig. 1D, and Fig. 1B-C, S1A-B), with a better inhibition of the accumulation of mRNAs transcribed from long segments (PB2, PB1 and PA) than short segments (M1, NS1). Moreover, the spliced M2 and NEP mRNAs were more impacted at both time points than their non-spliced counterparts, M1 and NS1, respectively (Fig. 1D).

Taken together, our data showed a stronger inhibition of the viral products obtained from long segments and from splicing as compared to unspliced and short segments. Nonetheless, unexpectedly high expression levels of IAV proteins could be detected in the presence of both MX1 proteins. This was particularly striking when looking at the potent inhibitions of viral production (Fig. 1A), and raised the possibility of blocks to infection occurring at later stages than viral RNA and protein synthesis.

### MX1 proteins perturb NP subcellular localization

We next wondered whether MX1 proteins could impact the localization of IAV proteins. A549 cells expressing FLAG-tagged Renilla (control), HsMX1 or MmMx1 were infected or not with A/Victoria/3/75 (MOI 1), fixed 8 h or 24 h later, and stained for FLAG and NP (Fig. 2A). First, we observed that the overall percentage of NP-expressing cells was only slightly decreased (by ∼7-15%) in the presence of HsMX1 or MmMx1, as compared to the control (Fig. 2A-B). This showed that NP expression was completely suppressed in only a minority of HsMX1- or MmMx1-expressing cells. In the control cells at 8 h post-infection, NP was found in three different localization patterns: uniquely localized in the nucleus (∼34% of the cells), both in the nucleus and the cytoplasm (∼58%) or uniquely in the cytoplasm (∼8%) (Fig. 2C). NP subcellular distribution largely resembled the control at 8 h post-infection in HsMX1-expressing cells. However, NP was predominantly found in the nuclei of MmMx1-expressing cells at this time point (∼75%) (Fig. 2A, C). At 24 h post-infection, as expected, NP was almost exclusively found in the cytoplasm of the control cells (∼83%) (Fig. 2A, C). In sharp contrast, NP was still exclusively found in the nucleus in the majority of MmMx1-expressing cells (∼67%), showing a dramatic impact of the murine antiviral protein on NP subcellular localization at this late time point (Fig. 2A, C). In HsMX1-expressing cells, NP was found in the three previous patterns, although at different proportions compared to the Renilla-expressing cells. Indeed, NP was uniquely present in the cytoplasm in ∼27% of HsMX1-expressing cells, present in both compartments in ∼63% of cells and still uniquely present in the nucleus in ∼10% of cells (compared to ∼83%, ∼16% and ∼1% of Renilla-expressing cells, respectively) (Fig. 2A, C). Similar data were obtained at 24 h post-infection with A/WSN/33 (Fig. S2A). Nucleocytoplasmic fractionations confirmed a strong decrease of NP accumulation in the cytoplasm of MmMx1-expressing cells at 24 h post-infection as compared to the control, while no difference could be observed for HsMX1 in this assay (Fig. S2B). Strikingly, however, at 24 h post-infection, in around ∼40% of NP-positive, HsMX1-expressing cells, NP was found accumulated into a dense, cytoplasmic structure localized in the perinuclear region (inserts from Fig. 2A and Fig. 2C for A/Victoria/3/75 and Fig. S2A for and A/WSN/33). Of note, HsMX1 was not found enriched in or around these structures at this time point (Fig. 2A and S2A).

**Figure 2.**
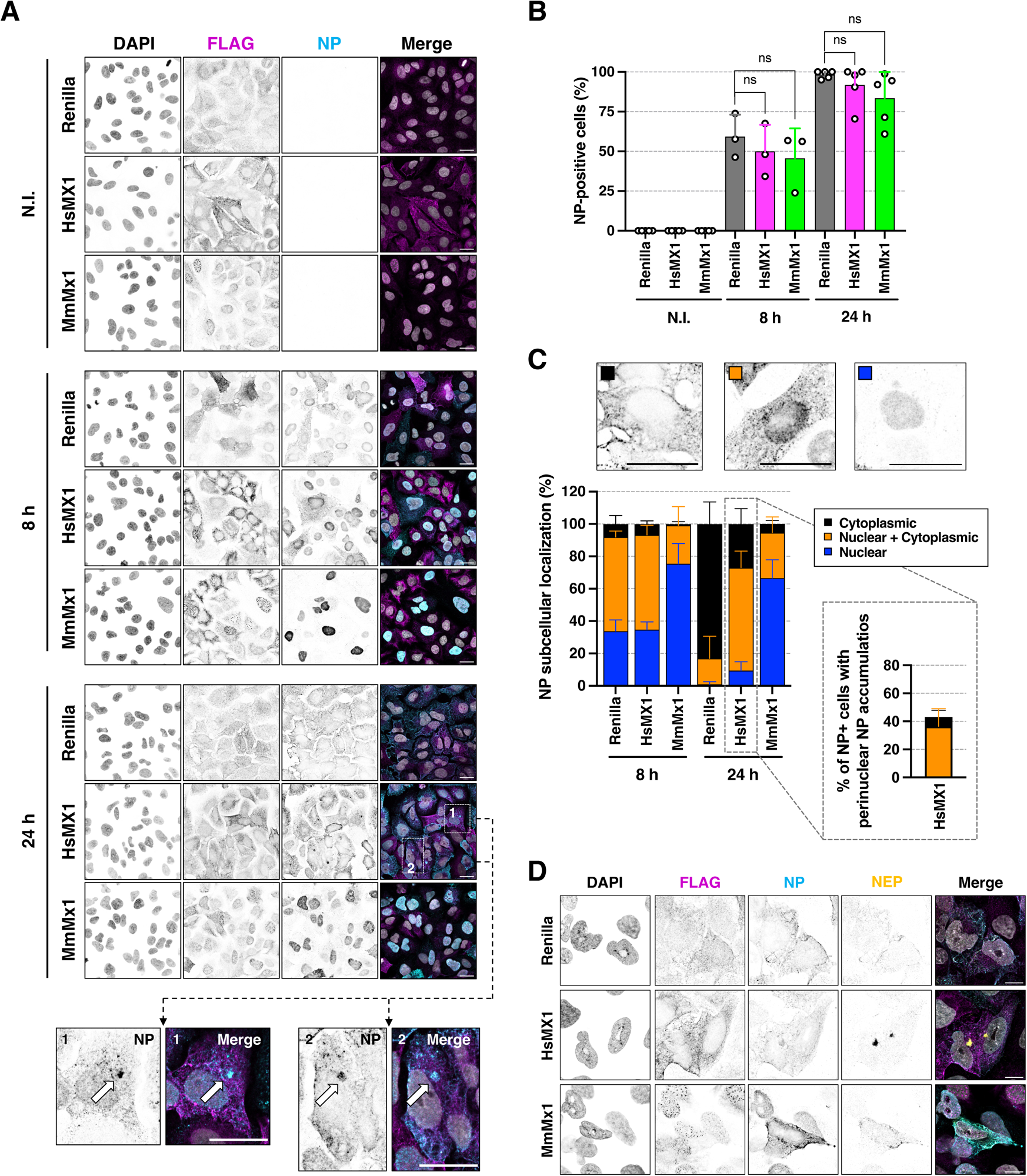
MX1 proteins impact NP subcellular localization at late timepoints post-infection. A549 cells expressing FLAG-tagged Renilla, HsMX1 or MmMx1 were infected or not (non-infected, N.I.) with A/Victoria/3/75 at MOI 1 for either 8 or 24 h. **A.** Representative confocal images of non-infected and infected cells are shown; FLAG (magenta), NP (cyan) and nuclei (grey). Single channels are shown as inverted greys. Insets 1 and 2: HsMX1-expressing cells infected for 24 h presenting a perinuclear NP accumulation. Scale bar: 20 µm. **B.** Quantification of the percentage of NP-positive cells in the different conditions. The mean and SD of 3 (8 h) or 5 (24 h) independent experiments are shown, with on average 266 cells analysed per condition. **C.** Top: representative images showing the different subcellular localization patterns of NP (exclusively cytoplasmic, both nuclear and cytoplasmic, and exclusively nuclear). Bottom: quantification of the percentage of cells showing each NP subcellular localization pattern at the indicated timepoints, with on average 370 cells in 5 fields analysed per condition, in 3 (8 h) and 5 (24 h) independent experiments. Insert: quantification of the percentage of NP-positive cells presenting the cytoplasmic NP accumulations (as shown with the arrows in inserts 1 and 2 from A) in the IAV-infected, HsMX1-expressing condition at 24 h post-infection. **D.** Representative Airyscan images of cells infected with A/Victoria/3/75 for 24 h; FLAG is represented in magenta, NP in cyan, NEP in yellow and nuclei in grey. Single channels are shown in inverted greys. Scale bar: 10 µm. Images are representative of 3 independent experiments.

Altogether, these data clearly showed that the subcellular distribution of NP was massively perturbed by MX1 proteins. An abnormal nuclear accumulation of NP, such as the one observed in the presence of MmMx1, could be reminiscent of viral nuclear export inhibition (Elton *et al*, 2001; Fournier *et al*, 2014; Guillon *et al*, 2022). As the Oxford Nanopore sequencing data showed significantly decreased levels of NEP mRNA in the presence of MmMx1 (Fig. 1D), we further analysed NEP expression and localization pattern by immunofluorescence (Fig. 2D and S2A; of note, the commercially available NEP antibodies are not suitable for immunoblot analyses). In the presence of MmMx1, NEP could not be detected in cells with nuclear NP and could only be detected in the rare cells presenting a cytoplasmic NP staining. This strongly suggested that NP abnormal nuclear accumulation might directly be connected to a defect in NEP expression and therefore nuclear export. Strikingly, in HsMX1-expressing cells, NEP accumulated in the same perinuclear structures as NP, at 24 h post-infection with both A/Victoria/3/75 and A/WSN/33 (Fig. 2D and S2A).

### HsMX1 induces an abnormal accumulation of IAV vRNPs in the vicinity of the MTOC

We sought to examine further the NP- and NEP-containing perinuclear structures observed in ∼40% of the NP-positive, HsMX1-expressing cells. As NEP is associated to trafficking vRNPs in the cytoplasm (O’Neill, 1998; Veler *et al*, 2022), we hypothesized that these structures could actually contain neo-synthesized vRNPs. Using FISH and immunofluorescence analyses, we observed that in addition to NP and NEP, M vRNA, PB2 and PA proteins were found in >99,6% of these perinuclear accumulations (Fig. 3A-C). This strongly pointed towards an accumulation of neo-synthesized vRNPs in these structures, which thereafter will therefore be referred to as ‘vRNP clusters’. Among other viral elements tested, M2 was found associated to all vRNP clusters whereas NS1 and M1 were only found in ∼8% and ∼35% of these structures, respectively (Fig. 3B-C, S2C and S3A-B). This argued against a general accumulation of viral elements in this area. Of note, these vRNP clusters were also detected in ∼40% of HsMX1-expressing HEK293T cells, showing that they were not cell-type specific (Fig. S3C-D).

**Figure 3.**
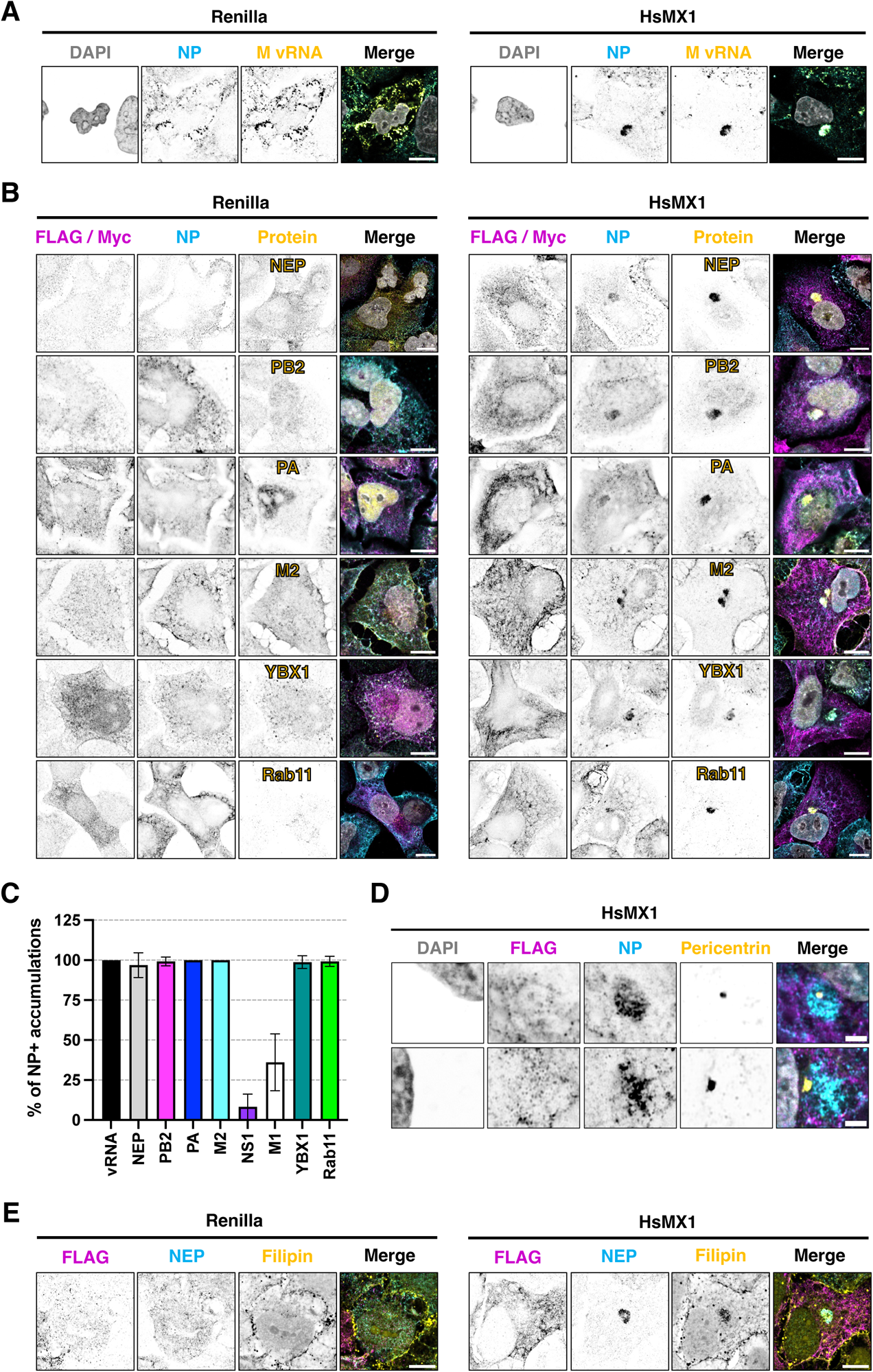
HsMX1 induces the sequestration of vRNPs and associated cellular factors in the vicinity of the MTOC. FLAG-tagged (or Myc-tagged, in the case of PA-FLAG staining) Renilla- or HsMX1-expressing cells were infected or not with A/Victoria/3/75 or A/Victoria/3/75-PA-FLAG at MOI 1, fixed 24 h later and stained as indicated. Single channels are shown in inverted greys. **A.** FISH coupled to immunofluorescence was performed and representative Airyscan images are shown with NP in cyan, M vRNA in yellow and nuclei in grey in the merged images. **B.** Representative Airyscan images of cells stained with the indicated markers. FLAG/Myc(-Renilla or -HsMX1) is shown in magenta, NP in cyan, the indicated proteins in yellow, and nuclei in grey. **C.** Quantification of the percentage of cells presenting NP accumulations containing the indicated markers on an average of 6 fields (with ∼100 cells) analysed per condition, in 3 (Rab11) or 2 (all the others) independent experiments. **D-E.** Representative Airyscan images of cells stained with the indicated markers. FLAG is shown in magenta, NP (**D**) or NEP (**E**) in cyan, Pericentrin (**D**) or Filipin (**E**) in yellow, and nuclei in grey. (**A-B**), (**D-E**) Representative images of 2 (M vRNA, PB2, PA-FLAG2X, Rab11, pericentrin) or 3 (M2, NEP, YBX1, and Filipin) independent experiments are shown. Scale bar: 10 µm (**A**, **B**, **E**) or 2 µm (**D**).

Next, we examined the localization of cellular proteins known to be involved in vRNP trafficking. HRB was not found in the HsMX1-induced vRNP clusters (Fig. S4A). This suggested that the dissociation of vRNPs from CRM1-Ran-GTP nuclear export complex was not perturbed in HsMX1-expressing cells. In contrast, both YBX1 and Rab11 were found in >99% of these structures (Fig. 3B-C). As IAV vRNPs are known to traffic towards the MTOC after nuclear export (Kawaguchi *et al*, 2012), we probed the MTOC localization with pericentrin and α-tubulin (Fig. 3D and S4B). We observed that the vRNP clusters were indeed found in the vicinity of the MTOC. Furthermore, staining with Filipin, a naturally fluorescent polyene macrolide binding specifically to sterols, revealed an accumulation of sterols within these structures (Fig. 3E). This strongly suggested that cholesterol-rich membranes were present, presumably under the form of Rab11a-positive vesicles (Kawaguchi *et al*, 2015). Such an observation might explain why M2 was detected in these structures, as it is known to associate to Rab11a-positive recycling endosomes (Rossman *et al*, 2010).

Transmission electron microscopy confirmed that infected HsMX1-expressing cells contained a peculiar perinuclear accumulation of vesicle-like structures of different sizes (Fig. 4A-B and S5A-B). These structures were never observed in the absence of infection or in Renilla-expressing cells, whether they were infected or not (Fig. 4A). Of note, the coalescence of vesicles observed in infected, HsMX1-expressing cells was sometimes found surrounded by the Golgi apparatus (Fig. S5B). Fluorescent imaging on fixed cells confirmed the proximity of the trans-Golgi network next to the vRNP clusters (Fig. S5C).

**Figure 4.**
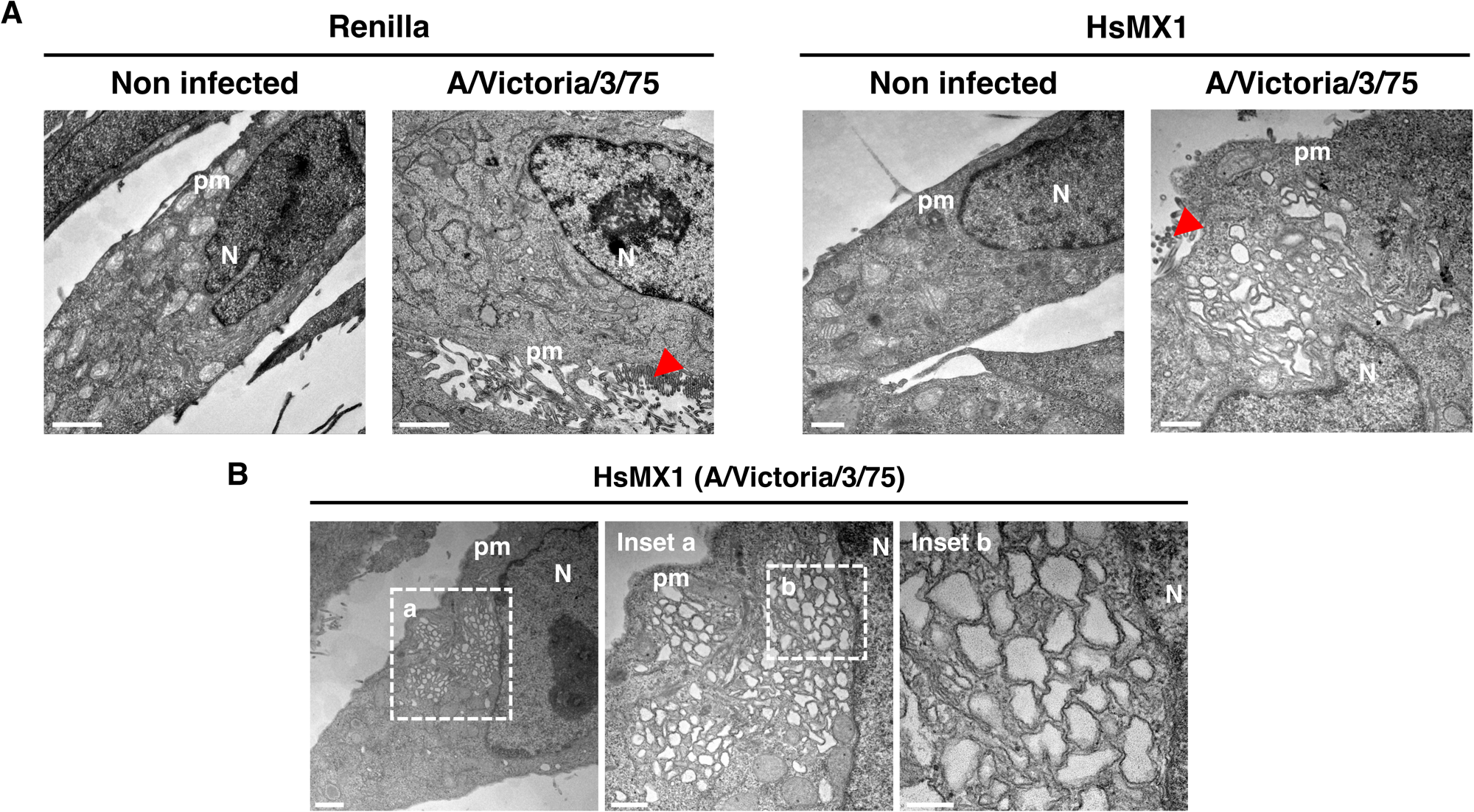
Ultrastructure of HsMX1-induced vRNP trafficking complex clusters. **A.** FLAG-tagged Renilla- and HsMX1-expressing cells were infected or not with A/Victoria/3/75 at MOI 1, fixed after 24 h and processed for transmission electron microscopy. Scale bar: 1 µm. An accumulation of vesicular membranes in proximity to the nucleus is shown inside the dashed lines in the HsMX1 condition. Scale bar: 1 µm (left) or 500 nm (right). **B.** A HsMX1-expressing, A/Victoria/3/75 infected cell presenting a perinuclear accumulation of vesicles polarized to one side of the nucleus with two insets shown at different zooms. Scale bar: 1 µm, inset a: 500 nm, inset b: 200 nm. (**A-B**) Representative images from 2 independent experiments are shown. N: nucleus, pm: plasma membrane. The red arrows indicate budding IAV particles.

Altogether these data showed that the nuclear export of YBX1-associated vRNPs and their association with Rab11-positive vesicles in the vicinity of the MTOC could occur in HsMX1-expressing cells. However, once in this area, the vRNPs abnormally accumulated in vesicular structures.

### HsMX1 severely impacts Rab11a-mediated vRNP trafficking in infected cells

To gain further insights into the formation of these HsMX1-induced, vRNP- and Rab11-containing structures, we performed live imaging analyses with cells stably co-expressing a GFP:Rab11a fusion along with either Renilla or HsMX1. In the absence of infection, GFP:Rab11a behaved similarly in A549-GFP:Rab11a-Renilla and A549-GFP:Rab11a-HsMX1 cells (Fig. 5A-B, Movies S1-2). Punctate structures, likely corresponding to Rab11a-positive endosomes, were observed, and could sometimes be seen trafficking to and from a perinuclear compartment, presumably corresponding to the endosome recycling compartment. In the context of IAV infection, however, drastic differences were observed (Fig. 5A-B, Movies S3-4). As previously described (Alenquer *et al*, 2019), in infected (Renilla-expressing) control cells, GFP:Rab11a was redistributed from a vesicular pattern to larger inclusions throughout the cytoplasm in >96% of the cells (Fig. 5A-B, Movie S3). In contrast, only ∼43 % of HsMX1-expressing, infected cells contained these large inclusions throughout the cytoplasm (Fig. 5A-B and Movie S4). In around half of the HsMX1-expressing cells which did not contain these large inclusions (i.e. in ∼25% of the total cells), GFP:Rab11a was actually found concentrated in a dense structure in the perinuclear region (Fig 5A-B), confirming with live imaging what was observed previously in fixed samples (Fig. 3B).

**Figure 5.**
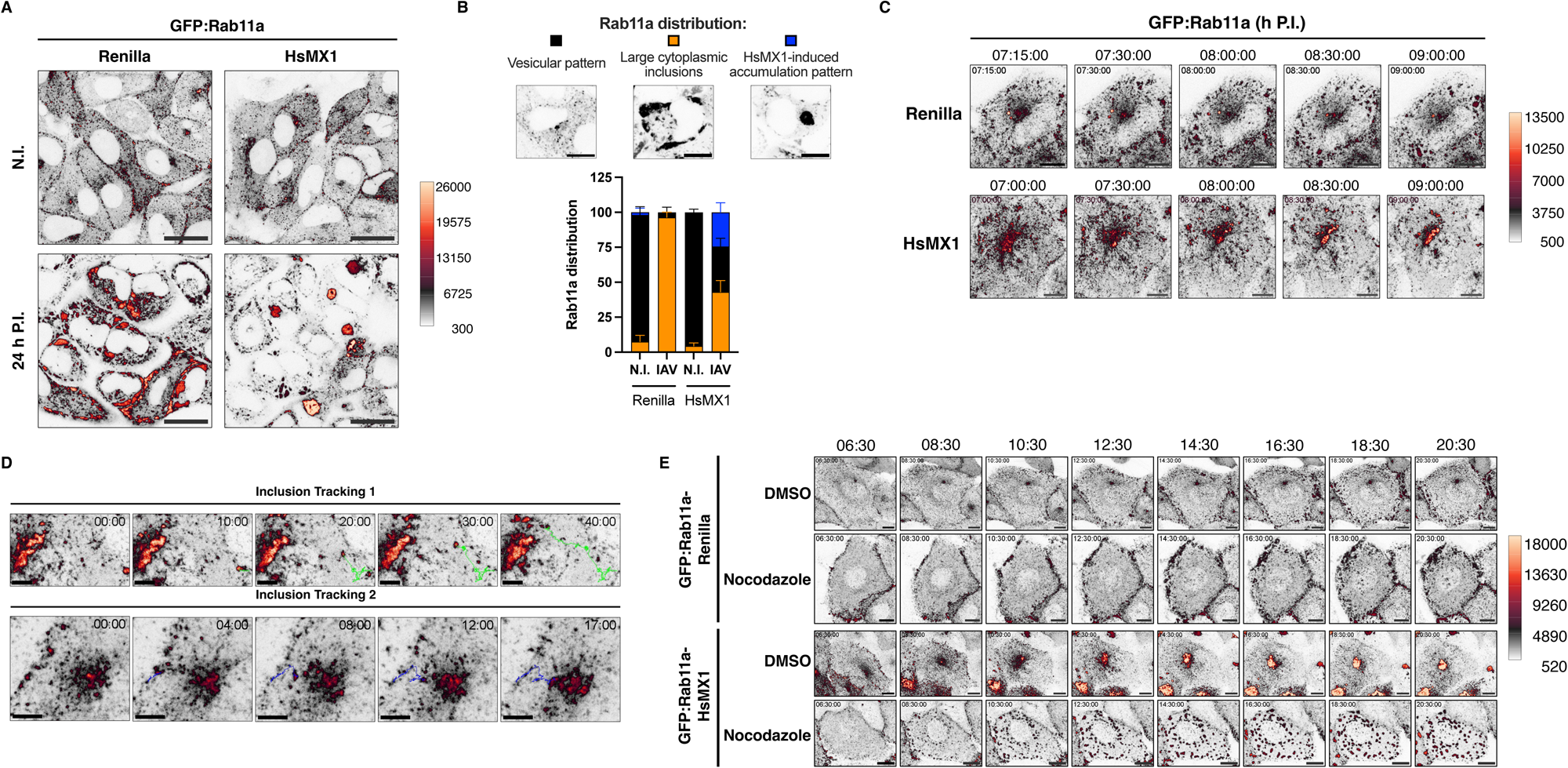
HsMX1 severely impacts Rab11a-mediated vRNP trafficking in infected cells. **A.** Representative stills of Movies S1-4 showing GFP:Rab11a in non-infected or in A/Victoria/3/75-infected, A549-GFP:Rab11a-Renilla or A549-GFP:Rab11a-HsMX1 cells, at 24 h post-infection. Scale bar: 20 µm. **B**. Top, representative images showing the different subcellular distribution patterns of Rab11a; bottom, quantification of the different patterns in 135 cells on average per condition, from Movies S1-4. **C.** Representative stills of maximum projected Z-stack acquisitions of GFP:Rab11a in A/Victoria/3/75-infected A549-GFP:Rab11a-Renilla and A549-GFP:Rab11a-HsMX1 cells. Images were recorded from ∼7 h post-infection (time points as indicated above the images), images correspond to Movies S5.2 and 6.1. Scale bar: 10 µm. **D.** The trajectory of two GFP:Rab11a inclusions from either Movie S6.1 (Inclusion Tracking 1; Movie S9) or Movie S6.2 (inclusion tracking 2; Movie S10) (i.e. A/Victoria/3/75-infected, A549-GFP:Rab11a-HsMX1 condition) were tracked manually and represented as green or blue lines, respectively. The time indicated represents the duration of the trafficking. Scale bar: 5µm. **E.** Time-course of A/Victoria/3/75-infected A549-GFP:Rab11a-Renilla or -HsMX1 cells treated with DMSO or Nocodazole from 3 h post-infection. Imaging was started at 6h30 post-infection and representative stills from Movies S11-12 and S13-14 taken every 2 h are shown, up to 20h30 post-infection. Scale bar: 10 µm. (**A, C, D, E**) Lookup table used is the MQ div-strawberry from the Neurocyto LUTs package on FIJI. Low intensities are white to black, medium intensities are from black to red and strong intensities are from red to yellow (of note, C and D use the same scale, shown in C). This representation allows easy visualization of weak and strong intensities on the same image.

We then set out to observe the formation of these structures in real time. Time course experiments in infected, Renilla-expressing control cells showed that GFP:Rab11a inclusions were found throughout the cytoplasm starting around 7 h post infection (Fig. 5C, Movies S5.1, 5.2, 5.3), as expected (Alenquer *et al*, 2019). In contrast, in infected HsMX1-expressing cells, a progressive and massive accumulation of GFP-Rab11a in a cytoplasmic area close to the nucleus, most likely in the vicinity of the MTOC, was observed from 7 h post-infection onwards (Fig. 5C, Movies S6.1, 6.2, 6.3). Of note, performing the live imaging experiments with a recombinant A/WSN/33 bearing a PA:mScarlet fusion showed that PA was present in these HsMX1-induced, GFP:Rab11a-containing perinuclear clusters and in the other cytoplasmic inclusions, strongly suggesting that they contained vRNPs as observed in fixed imaging (Movies S7-8, Fig. S6A and Fig. 3). Tracking some of the GFP:Rab11a inclusions from Movies S6.1-6.3 revealed that the trafficking process was not linear, with the GFP:Rab11a inclusions undergoing stop-and-start movements before reaching the perinuclear clusters (Fig. 5D, Movies S9-10). We therefore asked whether the microtubule network played a role in the formation of the large perinuclear clusters of Rab11a by evaluating the impact of nocodazole treatment on GFP:Rab11a dynamics in Renilla- or HsMX1-expressing A549 cells infected with A/Victoria/3/75. Nocodazole was added 3 h post-infection to prevent potential inhibitory effects on the initial phases of IAV replication (Banerjee *et al*, 2014). In the control cells, nocodazole induced an expected accumulation of GFP:Rab11a near the plasma membrane and a loss of the perinuclear localization pattern likely corresponding to the endosome recycling compartment (Fig. 5E and Movies S11-12). In HsMX1-expressing cells, the typical, large and unique perinuclear clusters of GFP:Rab11a were not observed in the presence of nocodazole (Fig. 5E, Movies S13-14). Instead, numerous and smaller GFP:Rab11a clusters were observed throughout the cytoplasm. These were not redirected towards the perinuclear region and remained relatively static over time, demonstrating a role for the microtubules in the formation of the HsMX1-induced, Rab11a-containing large perinuclear clusters. Immunofluorescence analyses confirmed that vRNP components NP and PB2 were found in the multiple cytoplasmic clusters, which were observed in ∼58% of infected cells at 24 h post-infection in the presence of nocodazole (Fig. 6A-B). Strikingly, HsMX1 was found to be associated to most of these vRNP clusters (∼68%) at 24 h post-infection, whereas it was absent at this time point in the DMSO condition (Fig. 6C and inset from Fig. 6A), as observed previously. Moreover, the multiple vRNP clusters observed with nocodazole were stained with Filipin, presumably indicating the presence of membranes (Fig. 6D). Altogether, this strongly suggested that these clusters, likely corresponding to HsMX1-Rab11a- and vRNP-associated vesicles, could be formed independently of a functional microtubule network. However, the latter was clearly necessary to traffic them back to and/or to maintain them in the perinuclear location in the vicinity of the MTOC.

**Figure 6.**
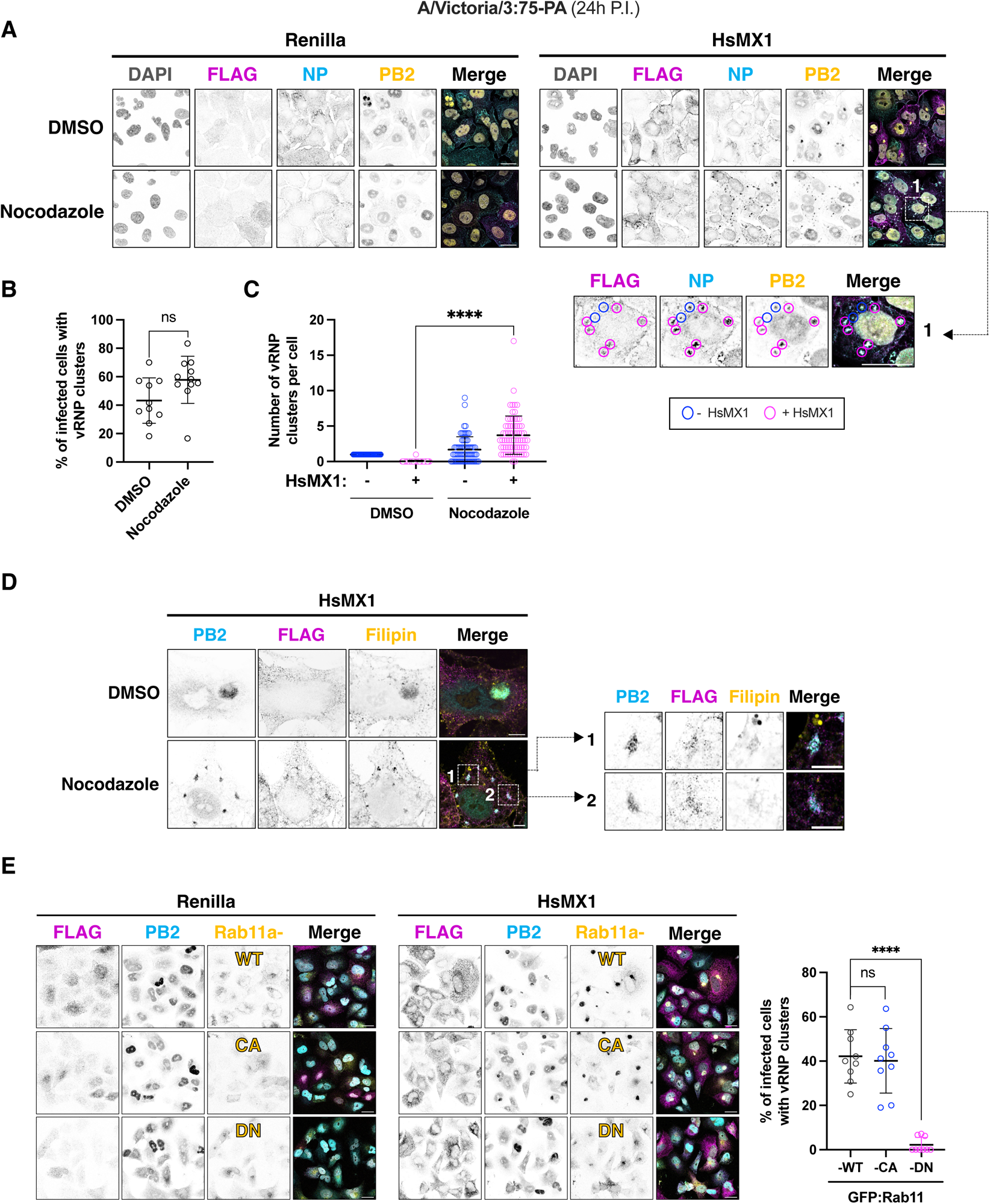
Microtubules play a role in the formation of HsMX1-induced, large vRNP clusters. **A.** A549 cells expressing FLAG-tagged Renilla or HsMX1 were infected with A/Victoria/3/75 at MOI 1, treated with DMSO or Nocodazole from 3 h post-infection and fixed at 24 h. Inset 1 shows a magnification of an HsMX1-expressing cell in the nocodazole condition, with magenta and blue circles indicating vRNP accumulations containing HsMX1 or not, respectively. FLAG is shown in magenta, NP in cyan, PB2 in yellow and nuclei in grey. Single channels are shown in inverted greys. Scale bar: 20 µm. **B.** Quantification of the percentage of HsMX1-expressing cells presenting vRNP clusters in the DMSO and nocodazole conditions **C.** Quantification of the number of vRNP clusters containing HsMX1 (Magenta symbols) or not (blue symbols) in the DMSO and nocodazole conditions. (**B**-**C**) 121 cells on average were counted from 10-11 fields per condition, in 2 independent experiments. **D.** Representative Airyscan images of A549 cells expressing FLAG-tagged HsMX1 were infected with A/Victoria/3/75 at MOI 1, treated with DMSO or Nocodazole from 3 h post-infection and fixed at 24 h post-infection. Insets 1 and 2 show magnified cytoplasmic accumulations in the Nocodazole condition. FLAG is shown in magenta, PB2 in cyan and Filipin in yellow. Single channels are represented in inverted greys. Scale bar: 5 µm. **E.** Left: Representative images of either FLAG-tagged Renilla- or HsMX1-expressing A549 cells, co-expressing GFP:Rab11a-WT, GFP:Rab11a-CA, or GFP:Rab11a-DN, as indicated, and infected with A/Victoria/3/75 at MOI 1 for 24 h. Single channels are shown in inverted greys, and FLAG is shown in magenta, PB2 in cyan, GFP-tagged Rab11a or mutants in yellow and nuclei in grey on the merged images. Scale bar: 20 µm. Right: quantification of the percentage of infected cells presenting vRNP accumulations (as evaluated here with PB2 staining) in the different conditions. 84 cells on average were counted on at least 9 fields per condition, in 2 independent experiments.

Next, we sought to explore the impact of Rab11a GTPase mutations on the HsMX1-induced, vRNP perinuclear clusters. To this aim, we used the Q70L constitutively active (CA) mutant, which is blocked in a GTP-bound form, and the S25N dominant negative (DN) mutant, blocked in a GDP-bound form (Ren *et al*, 1998). A549 cells stably expressing either Renilla or HsMX1, together with WT or mutant Rab11a fused to GFP, were infected by A/Victoria/3/75 (MOI 1) and fixed at 24 h post-infection for immunofluorescence analysis (Fig. 6E). Upon infection of HsMX1-expressing cells, GFP:Rab11a-WT and PB2 (used here as a surrogate for the vRNPs) were found in the perinuclear accumulations in ∼40% of cells, as observed previously. The GFP:Rab11a-CA mutant was similarly found in these accumulations together with PB2 in ∼40% of cells. However, in the case of GFP:Rab11a-DN, less than 2% of cells presented vRNP clusters, showing that the expression of this mutant prevented these accumulations from forming (Fig. 6E). Therefore, this argues that the presence of GTP-bound Rab11a and, presumably, the interaction of GTP-bound Rab11a with vRNPs, were pre-requirements for the vRNP clusters to be formed.

Taken together, these data showed that, while the presence of HsMX1 had no apparent impact on Rab11a behaviour in non-infected cells, it clearly affected the microtubule-dependant vesicular trafficking of Rab11a-associated vRNPs upon viral replication and induced their sequestration in the vicinity of the MTOC.

### HsMX1 transiently associates to Rab11a/vRNP trafficking complexes

As HsMX1 was found in most of the small vRNP clusters observed in the presence of nocodazole, we wondered whether it could transiently be associated to the trafficking vRNPs and/or to the vRNP large perinuclear clusters in the absence of the drug. To visualize HsMX1 in live imaging, we fused it to either GFP or Fluorescence-Activating and absorption-Shifting Tag (FAST, which has a smaller molecular weight than GFP (Plamont *et al*, 2016)). In addition to showing a cytoplasmic distribution not resembling that of WT HsMX1 (Fig. S6B and Movie S15), GFP:HsMX1 was poorly active against IAV in HEK293T cells (Fig. S6C). In contrast, FAST:HsMX1 showed a localization pattern highly resembling the classical honeycomb pattern of the WT protein (McKellar *et al*, 2023) (Fig. S6B and Movie S16). Moreover, FAST:HsMX1 retained antiviral activity in the same assay in HEK293T cells (Fig. S6C), as well as in stable A549 cells when co-expressed with WT HsMX1 (Fig. S6D). FAST:HsMX1 fusion protein was therefore selected for subsequent studies. In non-infected A549 cells, GFP:Rab11a and FAST:HsMX1 did not colocalize (Fig. 7A, left panel, Movies S17.1 and 17.2). In contrast, FAST:HsMX1 could be clearly detected colocalizing with the large Rab11a accumulations found in the perinuclear region at 9h30 post-infection (Fig. 7A, right panel, Movies S18.1, 18.2, 18.3). FAST-HsMX1 was also associated to cytoplasmic GFP:Rab11a inclusions trafficking back to the perinuclear accumulation (Fig. 7A-B, Movies S18.1, 18.2 and 18.3). However, in line with our immunofluorescence data, FAST:HsMX1 was no longer associated to these structures at 24 h post-infection (Fig. 7A, right panel, Movies S19.1, 19.2, 19.3). This suggested a transient association of HsMX1 to vRNP-Rab11a trafficking complexes, which was confirmed by imaging GFP:Rab11a and FAST:HsMX1 over the course of the infection (Fig. 7B and Movies S20.1, 20.2). FAST:HsMX1 association with GFP:Rab11a accumulations occurred from ∼7 h-7h30 to ∼10 h-10h30 post-infection and, after that, HsMX1 could not be detected anymore in these structures. Immunofluorescence on infected, HsMX1-expressing cells fixed at 9h30 post-infection confirmed the colocalization of HsMX1, NP and PB2 in the perinuclear region in ∼20% of infected cells, with HsMX1 detected in ∼42% of all vRNP clusters (Fig. 7C). The very transient nature of this colocalization between HsMX1 and viral proteins during the course of IAV infection most certainly explained why this was never reported before. Interestingly, this colocalization could potentially suggest a direct mechanism of action of HsMX1 on viral trafficking complexes.

**Figure 7.**
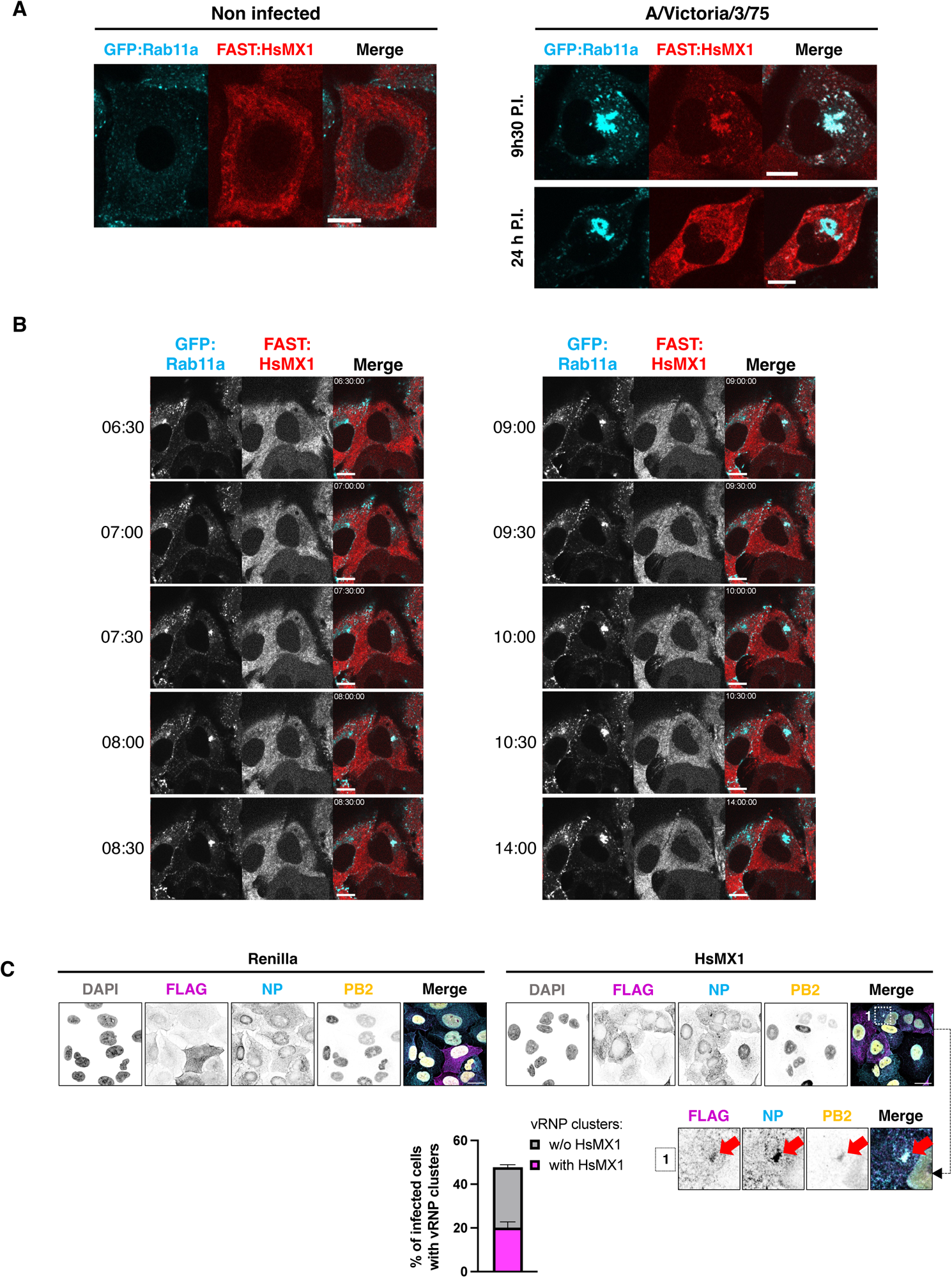
HsMX1 transiently associates to Rab11a/vRNP trafficking complex clusters. **A.** Representative stills from movies of A549-GFP:Rab11a-HsMX1-FAST:HsMX1 cells, non-infected or infected with A/Victoria/3/75 at MOI 1 for either 9h30 or 24 h (from Movies 17.1, 18.3 and 19.1). GFP:Rab11a is shown in cyan and FAST:HsMX1 in red. Scale bar: 10 µm. **B.** Time-course of A/Victoria/3/75-infected A549-GFP:Rab11a-HsMX1-FAST:HsMX1 cells. Representative stills from Movie 16.2 are shown and were taken every 30 minutes from 6h30 to 10h30 post-infection and at 14 h post-infection, as indicated. GFP:Rab11a is shown in cyan and FAST:HsMX1 in red. Single channels are represented in inverted grey. Scale bar: 10 µm. **C.** Representative confocal images of A549-FLAG-Renilla or A549-FLAG-HsMX1 cells infected with A/Victoria/3/75 at MOI 1 for 9h30. Inset 1 shows a magnified vRNP cluster in the HsMX1 condition (red arrow). FLAG is shown in magenta, NP in cyan PB2 in yellow and nuclei in grey. Single channels are shown in inverted greys. Scale bar: 20 µm. The graph represents the percentage of cells presenting vRNP clusters containing HsMX1 or not at 9h30 post-infection, as quantified on an average of 54 cells counted on 6 fields per condition, in 2 independent experiments. Results are representative of 2 (**C**) or 3 (**A-B**) independent experiments.

### HsMX1 requires dynein to induce the retrograde trafficking of IAV vRNPs and prevent viral production

Finally, we explored a potential role of cytoplasmic dynein, the molecular motor responsible for retrograde transport of a variety of cargos including Rab11a-recycling endosomes. To this aim, we used dynein pharmacological inhibition with the specific inhibitor Dynapyrazole A (Dyn-A) (Steinman *et al*, 2017). Dyn-A was added 3h post-infection to prevent any potential detrimental effects on the initial phases of IAV replication. The addition of Dyn-A at this time point in control cells had no impact on NP or PB2 localization patterns as observed by immunofluorescence 19h later (Fig. 8A). However, in HsMX1-expressing cells, the typical, large cytoplasmic vRNP clusters, observed in ∼52% of the cells in the DMSO conditions, were only found in ∼12% of the cells in the presence of Dyn-A. Moreover, there were no small vRNP clusters throughout the cytoplasm in the Dyn-A condition. Next, instead of adding the Dyn-A molecule early (3 h) post-infection, we tested its impact when added at 22 h post-infection, a time point at which the large perinuclear accumulations of HsMX1-induced vRNP clusters are established, and the cells were fixed for immunofluorescence analysis at 31 h post-infection (Fig. 8B). Nocodazole was used in parallel to Dyn-A. At this late time point of addition, nocodazole still induced the dispersal of the vRNP clusters throughout the cytoplasm similarly to when it was added early (Fig. 8B and 6A). In sharp contrast, when added at 22 h post-infection, Dyn-A did not affect the vRNP clusters (Fig. 8B). Taken together, these data showed that the microtubule network (or factors requiring it) was necessary for the formation and maintenance of the large vRNP clusters in the perinuclear region, whereas dynein activity seemed only necessary for their formation.

**Figure 8.**
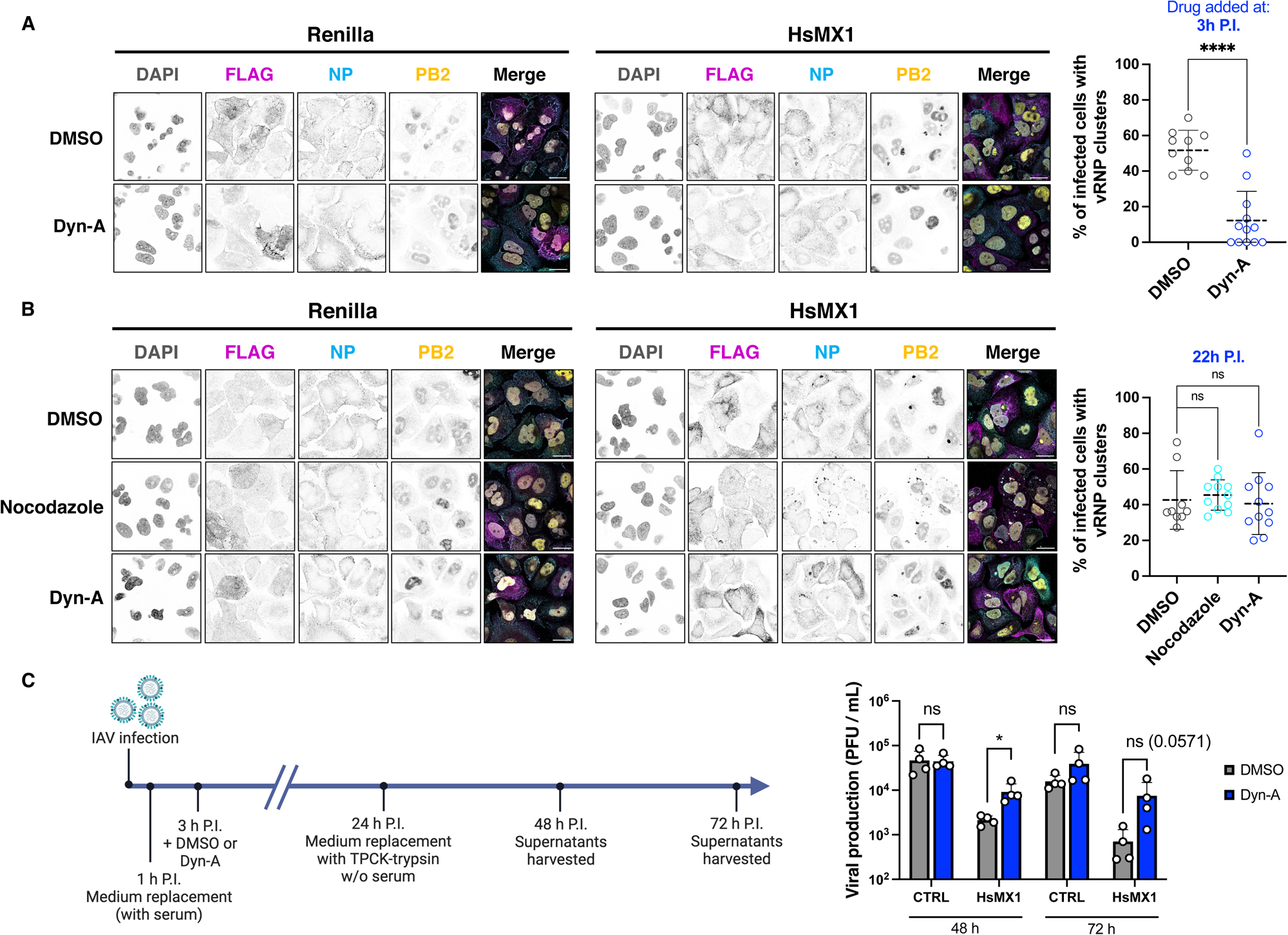
Dynein is involved in the formation of HsMX1-induced vRNP clusters. **A.** A549 cells expressing FLAG-tagged Renilla or HsMX1 were infected with A/Victoria/3/75 at MOI 1, treated with DMSO or Dynapyrazole-A (Dyn-A) at 3 h post-infection and stained for immunofluorescence analyses at 24 h by confocal microscopy. Left, representative images are shown (FLAG in magenta, NP in cyan, PB2 in yellow and nuclei in grey). Single channels are shown in inverted greys. Scale bar: 20µm. Right, quantification from A, with 82 cells on average counted from 12-14 fields per condition, in 2 independent experiments. **B.** A549 cells expressing FLAG-tagged Renilla or HsMX1 were infected with A/Victoria/3/75 at MOI 1, treated with DMSO, Nocodazole or Dynapyrazole-A at 22 h post-infection and fixed at 31 h post-infection for confocal imaging. FLAG is shown in magenta, NP in cyan, PB2 in yellow and nuclei in grey. Single channels are represented in inverted greys. Scale bar: 20 µm. Right, quantification from B, with 131 cells on average counted from 10 fields per condition, in 2 independent experiments. **C.** Same experiment as in A, but 24 h post infection the medium was replaced by serum-free, TPCK-trypsin medium (without drug) and the supernatants were harvested at 48 h and 72 h post-infection. Left, schematic of the experimental timeline; Right, viral production as measured by plaque assays. **A-C**. Results are representative of 2 (A-B) or 4 (C) independent experiments.

As Dyn-A strongly inhibited the formation of the large vRNP clusters, we then wondered whether it could rescue viral production in the presence of HsMX1. To assess this, we set up an experiment (depicted in Fig. 8C) in which the cells were infected and either DMSO or Dyn-A was added 3 h later in serum-containing medium, as above. However, in order to be able to assess infectious particle production, 24 h later, the medium was replaced with serum-free, TPCK-treated trypsin containing medium. The drug was not re-added at this time as we observed that Dyn-A precipitated in serum-free media, but its activity is known to be partially irreversible in serum-free conditions (Steinman *et al*, 2017), possibly providing a window to study its impact on infectious viral particle production. At 48 h and 72 h post-infection, the supernatants were harvested to measure infectious viral production in the different conditions by plaque assays (Fig. 8C). Dyn-A could partially and significantly rescue IAV infectious particle production in the presence of HsMX1 at 48 h post-infection, whereas it had no significant impact in control cells. The trend was similar at 72 h post-infection, despite not reaching statistical significance.

Taken together, these data support a model in which HsMX1 triggered the formation of abnormal Rab11a/vRNP clusters in the cytoplasm and favoured their dynein-dependant retrograde trafficking to a perinuclear region in the vicinity of the MTOC. Once there, HsMX1 dissociated from the vRNP clusters, which remained sequestered in this subcellular compartment.

## Discussion

Despite their discovery several decades ago, the detailed molecular mechanisms of action of MX1 proteins, both against IAV and in general, remain elusive. In this study, we revisited the inhibition of human IAV by MX1 proteins, using longer time points post-infection for our analyses than what was used before, and complementary techniques including fixed and live imaging. MmMx1 had previously been shown to strongly inhibit primary transcription from the longer IAV segments (Pavlovic *et al*, 1992). We indeed observed significantly decreased levels of IAV mRNAs by Oxford Nanopore sequencing, with the most potent effects on the mRNAs expressed from the longer IAV segments as well as from the spliced segments; of note, to our knowledge, the latter observation has never been reported before. However, we could surprisingly detect IAV protein expression in MmMx1-expressing cells, with notably only a minor impact on the percentage of cells expressing NP in comparison to the control cells. This apparent discrepancy with the previous data could be explained both by the timing of our experiments, analysed later than the initial ones (8 and 24 h herein versus 3-4 h post-infection (Pavlovic *et al*, 1992)) and the fact that we used human strains (versus avian strains, known to be more MX1-sensitive (Dittmann *et al*, 2008). We observed that NP abnormally accumulated in the nucleus of MmMx1-expressing cells during the late phases of viral replication, when it should be cytoplasmic. The decreased levels of NEP mRNA and protein (detected by Oxford Nanopore sequencing and immunofluorescence, respectively) pointed towards a reduced availability of NEP for nuclear export, which could explain this phenotype. Taking all these data into account, we can therefore hypothesize than MmMx1 does inhibit partially and/or delay primary transcription with more impact on the long and spliced segments, and this has multiple effects on IAV life cycle: i) a general decreased availability of viral proteins and in particular of those produced from the longer and spliced segments; and, consequently, ii) a decreased availability of NEP, probably leading to a profound defect in vRNP nuclear export. Overall, this leads to the potent replication inhibition of IAV observed in the presence of MmMx1. Of note, we were surprised to detect such high expression levels for viral proteins M1 or NS1 in MmMx1 expressing cells 24 h post-infection, despite a strong expression inhibition of viral polymerase subunits PB2 and PA. This suggested that i) primary transcription from segments 7 (M) and 8 (NS) was particularly efficient, in agreement with (Phan *et al*, 2021); and ii) the splicing machinery might not be efficiently recruited in these cells, leading to more unspliced viral mRNAs. Further studies will be needed to dissect the molecular mechanisms by which MmMx1 inhibits the accumulation of IAV mRNAs in general, and of spliced viral mRNAs in particular.

Contrary to MmMx1, HsMX1 had a minor impact on viral RNA and protein accumulation in infected cells, despite showing a robust phenotype in growth curve experiments. However, HsMX1 did impact, to some extent, the subcellular localization of NP, with some abnormal, exclusively nuclear localization at 24 h post-infection (in ∼10% of cells, vs ∼1% of the control cells) and an increased percentage of cells with both nuclear and cytoplasmic NP localization (in ∼64% of cells, vs ∼16% of the control cells). Whether the slight decrease of NEP mRNA levels with HsMX1 observed with Oxford Nanopore sequencing could explain this remains to be determined. Strikingly, however, in almost half of the cells in which the nuclear NP block was bypassed, HsMX1 did induce the peculiar accumulation of NP in the vicinity of the MTOC. Why this was not observed in all HsMX1-expressing cells in which NP nuclear export happened, and whether there might be a link with cell cycle progression for instance, remains to be determined. However, in addition to NP, viral proteins PB2, PA, NEP and M2, vRNA, cellular proteins YBX1, Rab11a as well as cholesterol-rich membranes were present in these perinuclear accumulations (of note, PB1 was not investigated). This strongly suggested that these accumulations corresponded to vRNPs associated to YBX1 and Rab11a-bearing vesicles, which did not traffic normally towards the plasma membrane in the presence of HsMX1. While there is evidence that NEP could remain associated to trafficking vRNPs (Veler *et al*, 2022), an association of M2 with vRNP transport has never been reported. We hypothesize that in the presence of HsMX1, M2-bearing, Rab11a-positive endosomes (Rossman *et al*, 2010) might be trafficked back to the perinuclear region, similarly to the ones with vRNPs, or their traffic could be prevented altogether. These hypotheses, involving Rab11a-vesicles bearing viral components, could be supported by an abnormal concentration of vesicles observed by EM in the perinuclear area 24h post infection, in infected, HsMX1-expressing cells. Another surprising observation was the fact than M1 was only associated to a fraction of the HsMX1-induced vRNP clusters. Indeed, M1 was suggested to remain attached to vRNPs trafficking towards the plasma membrane, in order to prevent reimport into the nucleus (Martin & Helenius, 1991; Whittaker *et al*, 1996; Babcock *et al*, 2004). However, a recent study, using GST pull-down with recombinant Rab11a and infected cell lysates, did not recover M1 together with Rab11a-associated vRNPs (Veler *et al*, 2022). Whether HsMX1 could induce an active exclusion of M1 from some of the trafficking vRNPs or whether M1 was often not present anymore at the time of HsMX1 action would remain to be determined.

The perinuclear sequestration of vRNPs induced by HsMX1 occurred subsequently to the interaction of vRNPs with GTP-bound Rab11a and was prevented by a dominant negative mutant of Rab11a. This showed that the trafficking step targeted could be concomitant to, or directly following the recruitment of Rab11a cofactor by vRNPs. Rab11a is a common host-dependency factor for various viruses in addition to IAV, including negative-strand RNA viruses such as Measles virus (MV), Sendai virus, Mumps virus or Respiratory Syncytial virus (RSV), or double stranded DNA Vaccinia virus (Nakatsu *et al*, 2013; Chambers & Takimoto, 2010; Hsiao *et al*, 2015; Katoh *et al*, 2015; Cosentino *et al*, 2022; Diot *et al*, 2023). Among these viruses only MV has so far been shown to be inhibited by HsMX1 (Schnorr *et al*, 1993; Schneider-Schaulies *et al*, 1994). It would be of interest to determine whether similar sequestration phenotypes are observed in the presence of HsMX1 for MV and other viruses utilizing Rab11a-mediated intracytoplasmic trafficking. Such investigations would assess whether the use of Rab11 is a common determinant for HsMX1 antiviral activity. Conversely, it might be interesting to evaluate the Rab11a-dependency of other viruses known to be inhibited by HsMX1. In a similar vein, perhaps one common determinant shared between certain viruses inhibited by HsMX1 is not a cellular cofactor *per se*, but the possibility to form cytoplasmic viral inclusions. Indeed, among the viruses listed above that utilize Rab11a, MV, Vaccinia virus and Respiratory Syncytial Virus (RSV) are known to form viral inclusions (Zhou *et al*, 2019; Kamahora *et al*, 1958; Rincheval *et al*, 2017). Looking beyond of Rab11a-dependency, other viruses have been reported to form viral inclusions (reviewed in (Etibor *et al*, 2021)) and among these viruses, Vesicular Stomatitis Virus (VSV), Rabies virus, Human Parainfluenza virus 3 and Ebola virus could be inhibited by HsMX1 (Heinrich *et al*, 2010; Nikolic *et al*, 2017; Zhang *et al*, 2013; Hoenen *et al*, 2012; Verhelst *et al*, 2013; Fuchs *et al*, 2017). It would therefore be interesting to determine if the capacity to form viral inclusions could be a factor denoting HsMX1 sensitivity.

While GFP:Rab11a behaviour seemed not to be impacted by HsMX1 expression in non-infected cells, we showed that, following IAV infection, sporadic cytoplasmic Rab11a inclusions were formed throughout the cytoplasm and abnormally trafficked in a retrograde manner towards the perinuclear region. HsMX1 was temporally associated with these structures, and used dynein and the microtubule network to induce this retrograde trafficking. IAV infection has been shown to induce a decrease of Rab11a binding to dynein through an unknown mechanism (Bhagwat *et al*, 2020). This could presumably promote the kinesin-mediated vRNP trafficking along microtubules towards the plasma membrane. It is therefore tempting to speculate that HsMX1 might be able to reverse this phenotype and reduce the kinesin-dependant transport of vRNPs, however preliminary experiments failed to confirm the initial findings in our experimental settings. HsMX1 is a mechanoenzyme belonging to the superfamily of Dynamin. Members of this superfamily regulate important biological functions by notably participating in various membrane remodelling events (Ramachandran & Schmid, 2018). These include endocytic vesicle fission and cytokinesis (e.g. dynamin), mitochondrial fusion (e.g. OPA1, mitofusins), mitochondrial and peroxisomal fission (e.g. Drp1), vacuolar dynamics and endosome regulation (Vps1). HsMX1 has never been shown to possess membrane remodeling activities in living cells, however it is known to possess lipid-tubulation abilities *in vitro* (Accola *et al*, 2002). In the light of our data, one might speculate that such an activity might be playing a role in the peculiar accumulation of vesicles observed in infected, HsMX1-expressing cells.

Remarkably, abnormal, perinuclear accumulations of IAV vRNPs have been previously reported in unrelated experimental conditions. Indeed, nucleozin, a pharmacological inhibitor inducing IAV NP aggregation, inhibits viral transcription and causes the subsequent cytoplasmic aggregation of vRNPs, along with Rab11a, in the perinuclear region (Kao *et al*, 2010; Su *et al*, 2010; Amorim *et al*, 2013). The main determinant for HsMX1 sensitivity has interestingly been reported to be NP and the current theorical model for HsMX1 anti-IAV activity involves potential physical interactions between NP and HsMX1 (Turan *et al*, 2004; Nigg & Pavlovic, 2015). It would be therefore tempting to speculate that, through interactions with NP, HsMX1 might induce some drastic changes in the vRNP structure, leading to similar consequences than NP aggregation by Nucleozin. Of note, however, physical interactions between HsMX1 and IAV NP are notoriously difficult to observe and have never been demonstrated in infected cells without the use of artificial conditions. Previous work used for instance dimeric mutants of HsMX1 with ectopic expression of IAV RdRp and NP in the context of a minireplicon assay, or HsMX1 fusion to a NLS (Turan *et al*, 2004; Nigg & Pavlovic, 2015). In our study, we provide the first evidence of a close proximity of HsMX1 with vRNP trafficking complexes during IAV infection, which could only be observed transiently. The transient nature of this association, observed both by live and fixed imaging in around ∼40% of cells, as well as the late timing, could explain why it had never been observed before. In our opinion, this transient association between HsMX1 and the vRNP-bearing, Rab11a-associated vesicles strongly suggests a direct mode of action of the antiviral protein against IAV vRNPs. Through structural modelling, HsMX1 has been proposed to form rings that might potentially surround viral components (Gao *et al*, 2011). Whether HsMX1 could sense and form these theorical “MX1 rings” around trafficking vRNP complexes in order to induce their retrograde traffic remains to be explored, through in depth cryo-EM studies for instance.

In conclusion, by combining infection experiments and extensive imaging analyses, we report a role of MX1 proteins in the late stages of IAV replication. Our study provides the first evidence of HsMX1 transiently associating with IAV vRNPs in infected cells and favouring dynein-dependant minus-end directed transport to inhibit viral trafficking towards the plasma membrane. These new observations should prompt to revisit the restriction phenotypes against the broad panel of RNA and DNA viruses inhibited by MX1 proteins in order to bring novel insights on the mechanisms of action of these broadly-active, antiviral dynamin-like large GTPases.

## Supporting information

Movie Legends and Link towards the movies

## Acknowledgments

We are thankful to María Moriel-Carretero, Arnaud Gauthier, Mikhail Baranov, Sandie Munier, Maïka Deffieu, Cécile Gauthier-Rouvière, Orestis Faklaris, Maria João Amorim, Magda Cannata-Serio and Jean-Baptiste Brault for sharing reagents and/or for very helpful discussions. We thank Kévin Terretaz for the development and sharing of the Visualization_toolset for FIJI used for image post-processing (https://github.com/kwolbachia/Visualization_toolset). We thank Nolwenn Jouvenet for critical reading of the manuscript.

This work was supported by the European Research Council (ERC) under the European Union’s Horizon 2020 research and innovation programme (grant agreement 759226, ANTIViR, to C. G.), the ATIP-Avenir program (to C.G.), the Institut National de la Santé et de la Recherche Médicale (INSERM) (to C.G.), a 3-year PhD studentship from the Ministry of Higher Education and Research (to J. M. and A.R.), a fourth year PhD funding from the Fondation pour la Recherche Médicale (FRM grant number [FDT202106013175] to J. M.), institutional funds from Centre National de la Recherche Scientifique (CNRS) and Montpellier University. O.M. and C.G. acknowledge support from the French research network on influenza viruses (ResaFlu; GDR2073) financed by the CNRS. Finally, the authors acknowledge the imaging facility MRI, a member of the national infrastructure France-BioImaging supported by the French National Research Agency (ANR-10-INBS-04).

## Author contributions

J.M., O.M. and C.G. designed the study, analysed the data and wrote the manuscript. J.M. carried out most of the experiments with assistance from MT (who did most molecular cloning and immunoblots), A.L.C.V. (who performed the Dyn-A rescue experiments and RT-qPCRs), M.A.A. (who performed immunoblots), A.R. (who produced viral stocks), O.M. (who produced viral stocks and performed the replication experiments). F.G.G. performed the Oxford Nanopore long read sequencing experiments with guidance from E.R. and E.L., and C.A. and S.M. analysed the Oxford Nanopore long read sequencing data. A.N. prepared the EM samples and acquired the EM images. N.N. provided stocks of the mScarlet reporter WSN virus. R.G. and B.D. provided essential reagents and guidance. All authors have read and approved the manuscript.

## Conflicts of interest statement

The authors have no conflicts of interest interests to declare in relation to this manuscript.

## Notes

### Competing Interest Statement

The authors have declared no competing interest.

### Summary of Updates

The "Movie Legends" file has been updated with the link to see the movies on FigShare and download them: https://figshare.com/articles/media/Movies_from_McKellar_et_al_bioRxiv_2024_Human_MX1_induces_the_cytoplasmic_sequestration_of_neo-synthesized_influenza_A_virus_vRNPs/25289647 This link has also been added in the Methods section of the manuscript file. The title of the manuscript has been modified ('induces' has been replaced by 'orchestrates')

