## Supplementary material for "Human MX1 orchestrates the cytoplasmic sequestration of neo-synthesized influenza A virus vRNPs": Legends of supplementary Figures

**Figure S1. A.** A549 cells expressing Myc-tagged Renilla, HsMX1 or MmMx1 were infected or not (N.I.) with A/Victoria/3/75-PA-FLAG at MOI 1 for 24 h, lysed and viral protein accumulation was analysed by immunoblot; Actin served as a loading control. Representative immunoblots (out of 2 independent experiments) are shown. **B.** A549 cells stably expressing FLAG-tagged Renilla, -HsMX1 or -MmMx1 were infected with A/Victoria/3/75-PA-Nanoluciferase at MOI 1 for 24 h and relative infection efficiencies were determined by monitoring Nanoluciferase activity. The graph represents the mean of 5 independent biological replicates. **C.** RT-qPCR analysis of PA (left) and M1 (right) mRNAs from A549 cells expressing FLAG-tagged CTRL (Renilla), HsMX1 or MmMx1 and infected or not with A/Victoria/3/75 at MOI 1 for 8 or 24 h. The mean and SD of 3 independent experiments are shown.

**Figure S2. A.** Representative confocal images of A549 cells stably expressing FLAG-tagged Renilla, HsMX1 or MmMx1 infected with A/WSN/33 at MOI 1 for 24 h. FLAG is represented in magenta, NP in cyan, NEP in yellow and nuclei in greys. Single channels are shown in inverted grey. Scale bar: 20 µm. Right panel, quantification of the percentage of cells showing a specific NP subcellular localization (top), with on average 400 cells analysed in 2 independent experiments, and quantification of the percentage of NP-positive cells presenting NP large accumulations (as shown in inserts from Fig. 2A and in Fig. 3A-B) in the IAV-infected, HsMX1-expressing cells (bottom). **B.** Nucleocytoplasmic fractionation of FLAG-tagged Renilla-, HsMX1- or MmMx1-expressing A549 cells infected or not with A/Victoria/3/75 at MOI 1 for 24 h. GAPDH and Histone 2B (H2B) were used as controls for cytoplasmic and nuclear fraction purity, respectively. Representative immunoblots (out of 2 independent experiments) are shown. W: Whole cell lysate, C: Cytoplasmic fraction, N: Nuclear fraction. **C.** Representative confocal images of A549 cells stably expressing FLAG-tagged Renilla, HsMX1 or MmMx1 infected with A/Victoria/3/75 at MOI 1 for 24 h. Single channels are shown in inverted grey, and on the merge, FLAG is shown in magenta, NP in cyan, NS1 in yellow and nuclei in greys. Scale bars: 20 µm. (**A,C**). Images are representative of 2 independent experiments.

**Figure S3. A.** Representative confocal images of A549 cells stably expressing FLAG-tagged Renilla, HsMX1 or MmMx1 infected with A/Victoria/3/75 at MOI 1 for 24 h. FLAG is represented in magenta, NP in cyan, M1 in yellow and nuclei in grey. Single channels are shown in inverted greys. Scale bars: 20 µm. **B.** Representative Airyscan images of FLAG-HsMX1-expressing A549 cells infected with A/Victoria/3/75 at MOI 1 for 24 h: example of vRNP accumulations (red arrows) without M1 (1) or containing M1 (2) (same colour code as in A), as quantified in Fig. 2C. Scale bars: 10 µm. **C.** Representative Airyscan images of HEK293T cells stably expressing FLAG-tagged Renilla or HsMX1 infected with A/Victoria/3/75 at MOI 1 for 24 h. Single channels are shown in inverted greys, and on the merge, FLAG is represented in magenta, NP in cyan, PB2 in yellow and nuclei in grey. Scale bars: 10 µm. **D.** Quantification of the percentage of NP-positive cells presenting vRNP accumulations (as quantified with NP and PB2 staining) in the IAV-infected, HsMX1-expressing HEK293T cells, with on average 70 cells analysed per condition (each symbol represents the % of positive cells per field). (**A**-**D**) Images and data are representative of 2 independent experiments.

**Figure S4.** Representative confocal images of A549 cells stably expressing FLAG-tagged Renilla or HsMX1 infected or not with A/Victoria/3/75 at MOI 1 for 24 h. Single channels are shown in inverted greys and on the merged images, FLAG is represented in magenta, NP in cyan, HRB (**A**) or α-Tubulin (**B**) in yellow and nuclei in grey. Scale bars: 20 µm (**A**) and 2 µm (**B**). Images are representative of 2 (**A**) or 3 (**B**) independent experiments. (**A**) Right panel: quantification of the percentage of NP-positive accumulations containing HRB with on average 51 cells analysed in 2 independent experiments.

**Figure S5. A-B.** Representative EM images of HsMX1-expressing cells infected with A/Victoria/3/75 at MOI 1 for 24 h. Scale bar: 500 nm (**A**), 200 nm (**B**). N: nucleus, pm: plasma membrane, g: golgi. The red arrows indicate budding IAV particles. **C.** Representative Airyscan images of A549 cells stably expressing FLAG-tagged HsMX1 and infected with A/Victoria/3/75 at MOI 1 for 24 h. Single channels are shown in inverted greys and on the merged images, FLAG is represented in magenta, NP in cyan, TGN46 in yellow and nuclei in grey. Scale bars: 10 µm. (**A-C**) Images are representative of 2 independent experiments.

**Figure S6. A.** Representative stills of Movies S7-8 showing GFP:Rab11a and PA-mScarlet in **A/WSN/33**-PA-mScarlet-infected, A549-GFP:Rab11a-Renilla or A549-GFP:Rab11a-HsMX1 cells, respectively, at 12 h post-infection. Scale bar: 10 µm. **B.** A549 cells stably expressing GFP:HsMX1, or FAST:HsMX1 along with FLAG-HsMX1, as indicated, were submitted to live imaging. Snapshots from Movies S15-16 are shown in inverted greys. Scale bar: 10 µm. Images are representative of 2 independent experiments. **C.** Infection reporter assay of HEK293T cells transfected with Renilla (for normalization purposes) and either an empty vector (CTRL) or wild-type, GFP-tagged or FAST-tagged HsMX1 constructs as well as with a polI-driven, Firefly expressing IAV minigenome, as in (McKellar *et al*, 2023). Infection with A/Victoria/3/75 was performed at MOI 1 for 24 h, cells were lysed and Firefly luciferase levels were measured and normalized to Renilla luciferase levels. The graph represents the mean and SD of 3 independent biological replicates. **D.** A549 cells stably expressing wild-type HsMX1 (WT), FAST:HsMX1, or both FAST:HsMX1 and FLAG-HsMX1 (FAST/WT) were infected with A/Victoria/3/75-NLuc at MOI 1 for 16 h and relative infection efficiencies were determined. The graph represents the mean of 2 independent biological replicates.
