## Supplementary figures and images for "Human MX1 orchestrates the cytoplasmic sequestration of neo-synthesized influenza A virus vRNPs"

### Figure S1

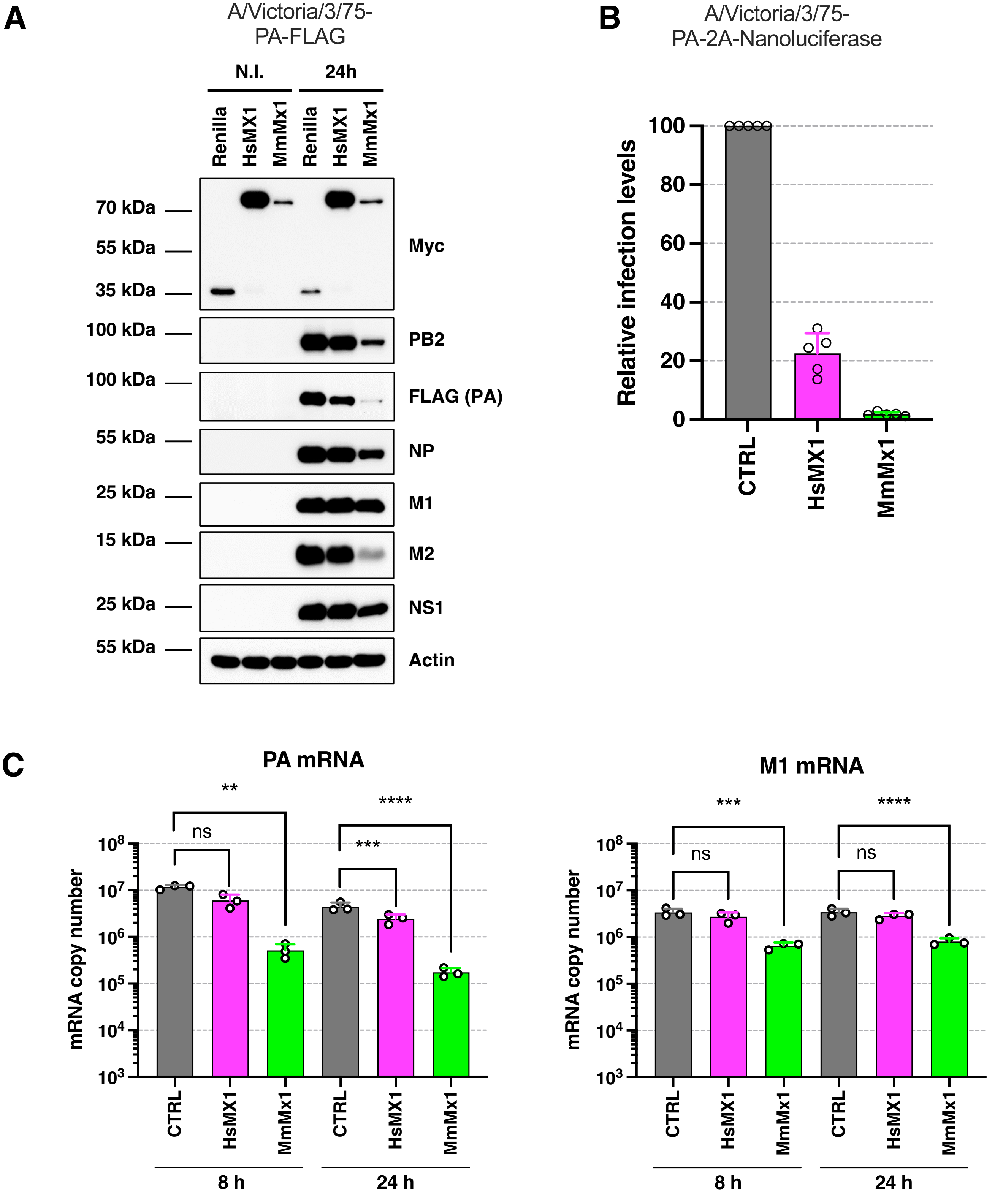

### Figure S2

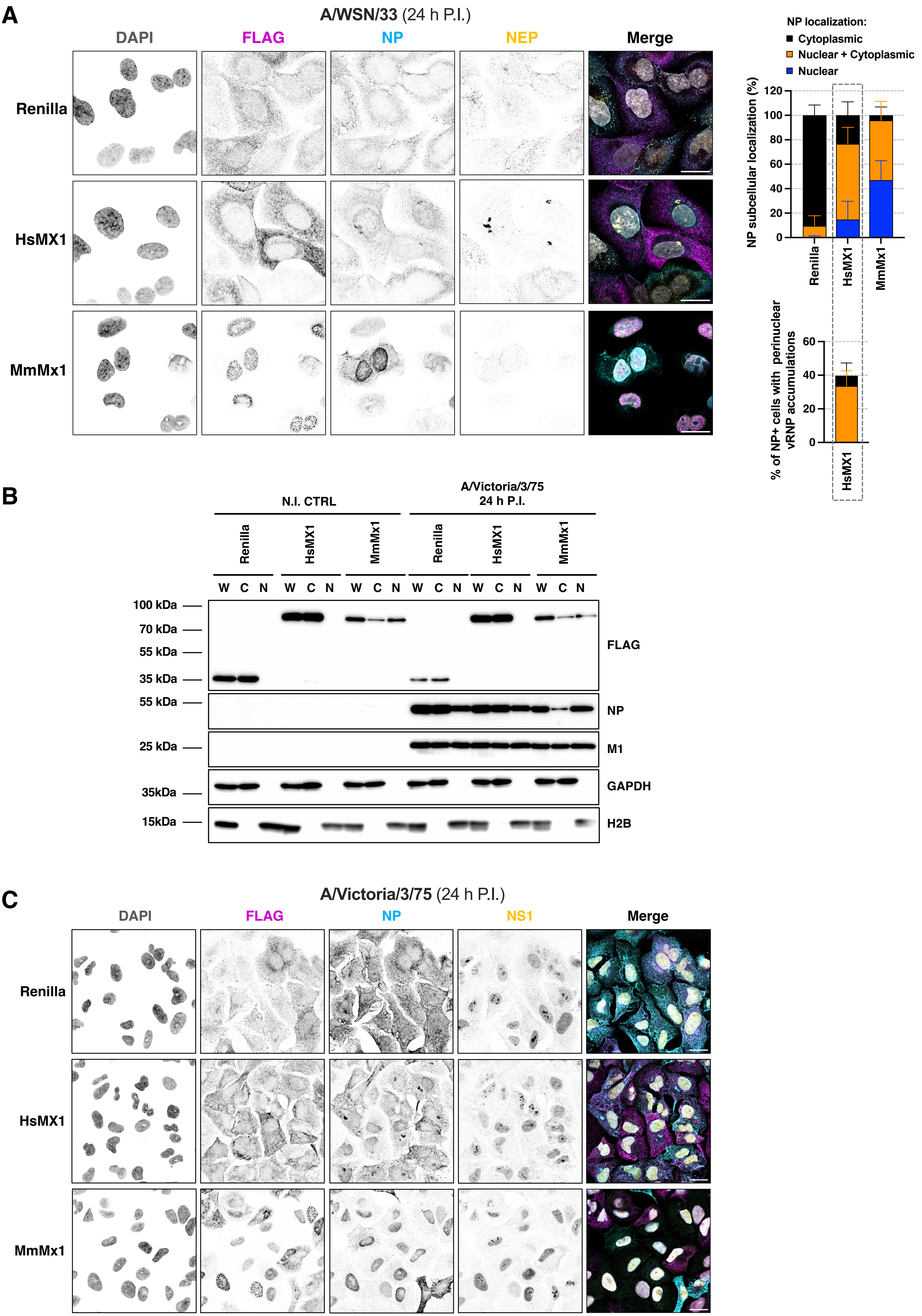

### Figure S3

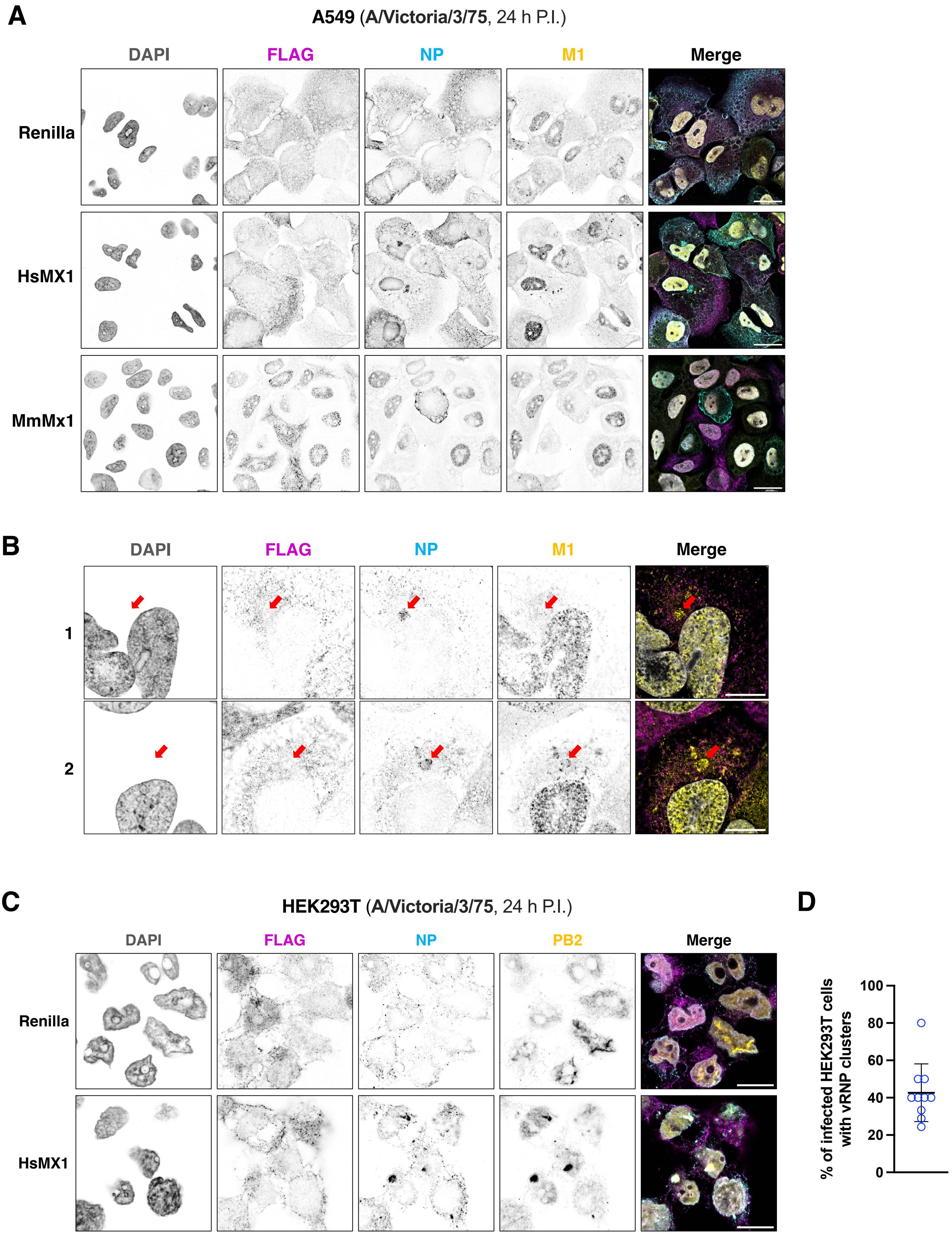

### Figure S4

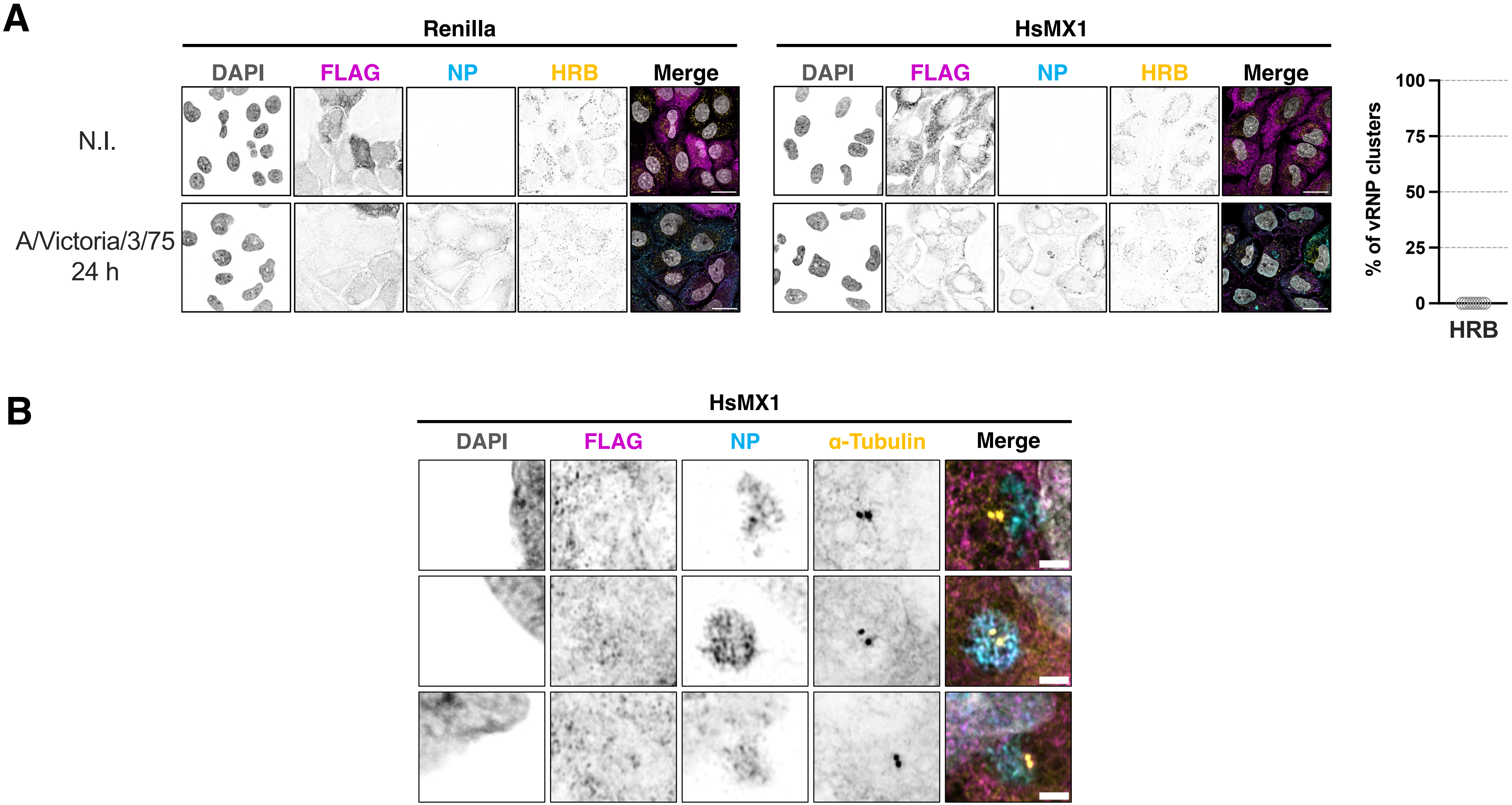

### Figure S5

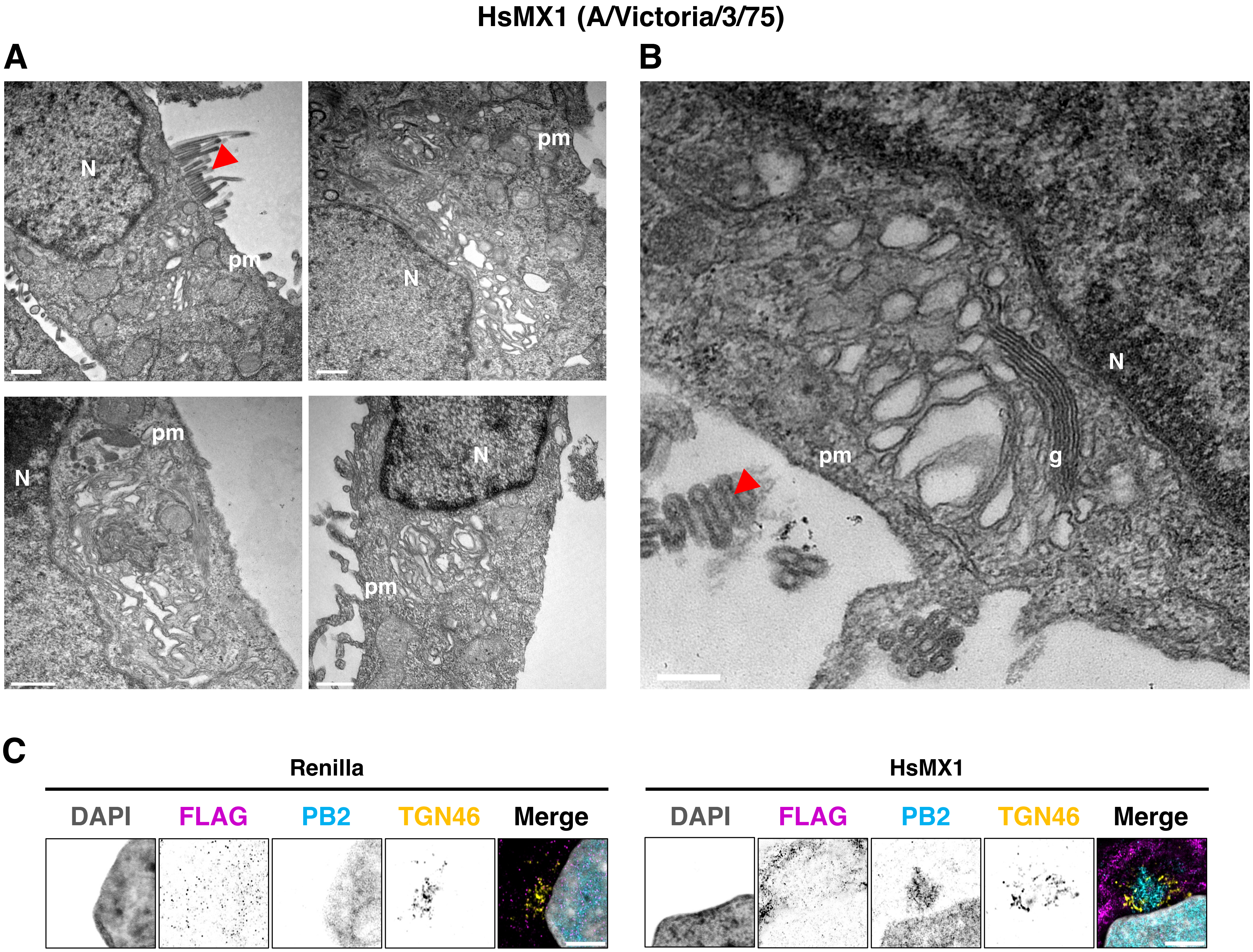

### Figure S6

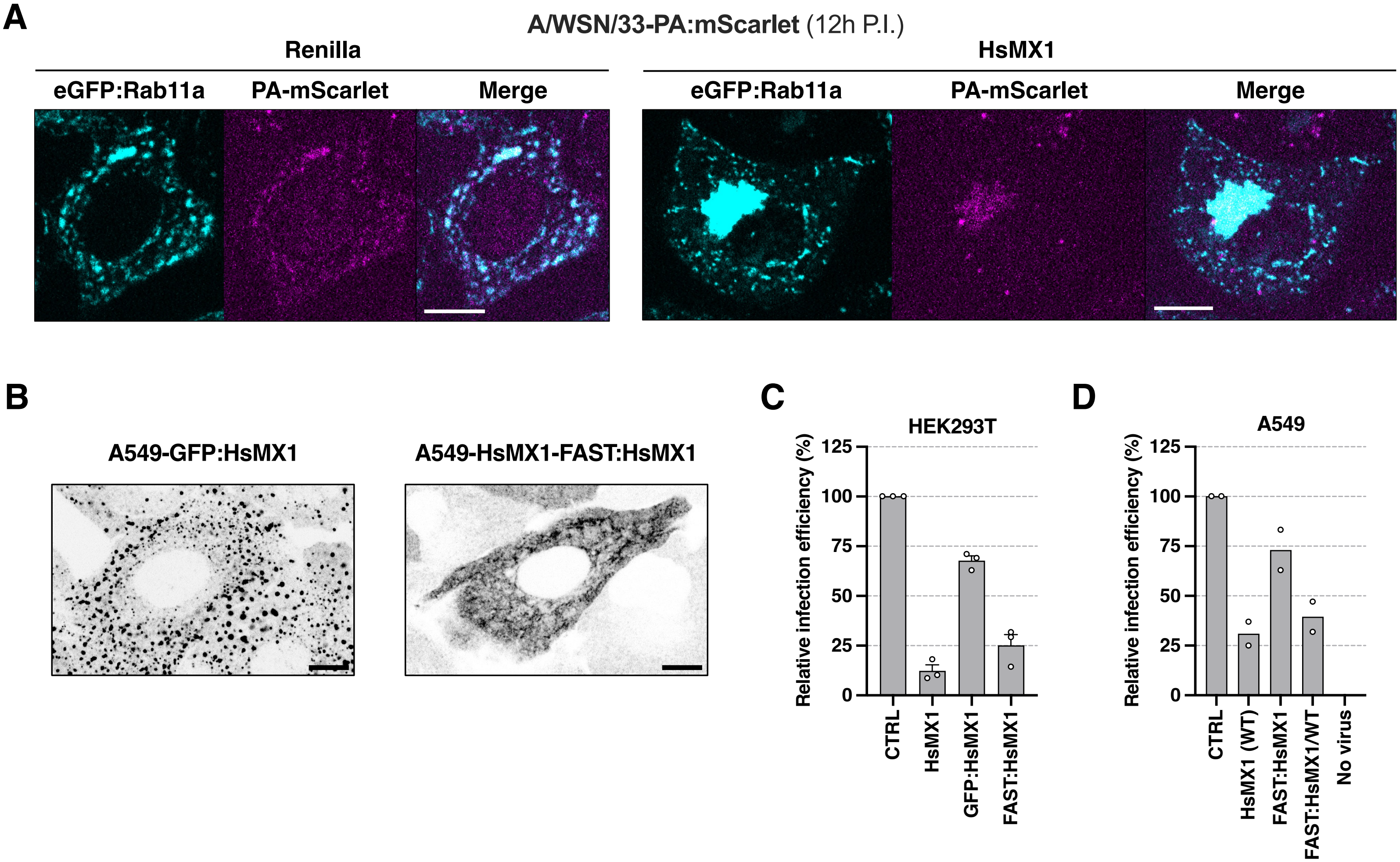
