## Supplementary material for "Human MX1 orchestrates the cytoplasmic sequestration of neo-synthesized influenza A virus vRNPs": Movie Legends and Link towards the movies

The Movies can be seen on (and downloaded from) [FigShare](https://figshare.com/articles/media/Movies_from_McKellar_et_al_bioRxiv_2024_Human_MX1_induces_the_cytoplasmic_sequestration_of_neo-synthesized_influenza_A_virus_vRNPs/25289647):

<https://figshare.com/articles/media/Movies_from_McKellar_et_al_bioRxiv_2024_Human_MX1_induces_the_cytoplasmic_sequestration_of_neo-synthesized_influenza_A_virus_vRNPs/25289647>

**Movie S1:** A549-GFP:Rab11a-Renilla-expressing cells in non-infected conditions. A single Z-slice is represented**.** GFP:Rab11a is represented in the MQ div-strawberry LUT. Low intensities are white to black, medium intensities are from black to red and strong intensities are from red to yellow. Movie representative of three independent experiments. One image per 250 ms for 1 minute. Scale bar: 20 µm.

**Movie S2:** A549-GFP:Rab11a-HsMX1-expressing cells in non-infected conditions. A single Z-slice is represented**.** GFP:Rab11a is represented in the MQ div-strawberry LUT. Low intensities are white to black, medium intensities are from black to red and strong intensities are from red to yellow. Movie representative of three independent experiments. One image per 250 ms for 1 minute. Scale bar: 20 µm.

**Movie S3:** A549-GFP:Rab11a-Renilla-expressing cells in A/Victoria/3/75 infected conditions 24 h post-infection at MOI 1. A single Z-slice is represented**.** GFP:Rab11a is represented in the MQ div-strawberry LUT. Low intensities are white to black, medium intensities are from black to red and strong intensities are from red to yellow. Movie representative of three independent experiments. One image per 250 ms for 1 minute. Scale bar: 20 µm.

**Movie S4:** A549-GFP:Rab11a-HsMX1-expressing cells in A/Victoria/3/75 infected conditions 24 h post-infection at MOI 1. A single Z-slice is represented**.** GFP:Rab11a is represented in the MQ div-strawberry LUT. Low intensities are white to black, medium intensities are from black to red and strong intensities are from red to yellow. Movie representative of three independent experiments. One image per 250 ms for 1 minute. Scale bar: 20 µm.

**Movie S5.1-5.3:** A549-GFP:Rab11a-Renilla-expressing cells infected with A/Victoria/3/75. Image acquisition was started 7h15 minutes post-infection, every 6 seconds until the end of the respective films. Films are represented as maximum Z-projections**.** GFP:Rab11a is represented in the MQ div-strawberry LUT. Low intensities are white to black, medium intensities are from black to red and strong intensities are from red to yellow. The movies are representative of three independent experiments. Scale bar: 10 µm.

**Movie S6.1-6.3:** A549-GFP:Rab11a-HsMX1-expressing cells infected with A/Victoria/3/75. Image acquisition was started 7 h post-infection, every 3 (6.1, 6.2) or 6 (6.3) seconds until the end of the respective films. Films are represented as maximum Z-projections**.** GFP:Rab11a is represented in the MQ div-strawberry LUT. Low intensities are white to black, medium intensities are from black to red and strong intensities are from red to yellow. The movies are representative of three independent experiments. Scale bar: 10 µm.

**Movie S7:** A549-GFP:Rab11a-Renilla-expressing cells infected with WSN-PA:mScarlet for 12 h at MOI 1. One image was acquired every 2 seconds for 30 seconds. GFP:Rab11a is represented in cyan and PA:mScarlet in magenta. The movie is representative of three independent experiments. Scale bar: 10 µm.

**Movie S8:** A549-GFP:Rab11a-HsMX1-expressing cells infected with WSN-PA:mScarlet for 12 h at MOI 1. One image was acquired every 2 seconds for 30 seconds. GFP:Rab11a is represented in cyan and PA:mScarlet in magenta. The movie is representative of three independent experiments. Scale bar: 10 µm.

**Movie S9:** Inset of Movie S6.1. A549-GFP:Rab11a-HsMX1-expressing cells infected with A/Victoria/3/75. Image acquisition was started 7 h post-infection, every 3 seconds until the end of the respective films. Films are represented as maximum Z-projections**.** GFP:Rab11a is represented in the MQ div-strawberry LUT. Low intensities are white to black, medium intensities are from black to red and strong intensities are from red to yellow. Scale bar: 5 µm.

**Movie S10:** Inset of Movie S6.2. A549-GFP:Rab11a-HsMX1-expressing cells infected with A/Victoria/3/75. Image acquisition was started 7 h post-infection, every 6 seconds until the end of the respective films. Films are represented as maximum Z-projections**.** GFP:Rab11a is represented in the MQ div-strawberry LUT. Low intensities are white to black, medium intensities are from black to red and strong intensities are from red to yellow. Scale bar: 5 µm.

**Movie S11:** DMSO-treated A549-GFP:Rab11a-Renilla-expressing cells infected with A/Victoria/3/75 for 6h30 at MOI 1 and imaged until 21h25 post-infection. Representation is a maximum Z-projection**.** GFP:Rab11a is represented in the MQ div-strawberry LUT. Low intensities are white to black, medium intensities are from black to red and strong intensities are from red to yellow. The movie is representative of three independent experiments. Scale bar: 10 µm.

**Movie S12:** Nocodazole-treated A549-GFP:Rab11a-Renilla cells infected with A/Victoria/3/75 for 6h30 at MOI 1 and imaged until 21h25 post-infection. Representation is a maximum Z-projection**.** GFP:Rab11a is represented in the MQ div-strawberry LUT. Low intensities are white to black, medium intensities are from black to red and strong intensities are from red to yellow. The movie is representative of three independent experiments. Scale bar: 10 µm.

**Movie S13:** DMSO-treated A549-GFP:Rab11a-HsMX1 cells infected with A/Victoria/3/75 for 6h30 at MOI 1 and imaged until 21h25 post-infection. Representation is a maximum Z-projection**.** GFP:Rab11a is represented in the MQ div-strawberry LUT. Low intensities are white to black, medium intensities are from black to red and strong intensities are from red to yellow. The movie is representative of three independent experiments. Scale bar: 10 µm.

**Movie S14:** Nocodazole-treated A549-GFP:Rab11a-HsMX1 cells infected with A/Victoria/3/75 for 6h30 at MOI 1 and imaged until 21h25 post-infection. Representation is a maximumZz-projection**.** GFP:Rab11a is represented in the MQ div-strawberry LUT. Low intensities are white to black, medium intensities are from black to red and strong intensities are from red to yellow. The movie is representative of three independent experiments. Scale bar: 10 µm.

**Movie S15:** A549-GFP:HsMX1-expressing cell. One image was acquired every 500 ms for 60 seconds. GFP:HsMX1 is represented in inverted grey. The movie is representative of three independent experiments. Scale bar: 10 µm.

**Movie S16:** A549-FAST:HsMX1-expressing cell. One image was acquired every 500 ms for 60 seconds. FAST:HsMX1 is represented in inverted grey. The movie is representative of three independent experiments. Scale bar: 10 µm.

**Movie S17.1-17.2:** Non-infected A549-GFP:Rab11a-FAST:HsMX1-expressing cell. One image was acquired every 1 second for 30 seconds. GFP:Rab11 is represented in cyan and FAST:HsMX1 in red. The movies are representative of three independent experiments. Scale bar: 10 µm.

**Movie S18.1-18.3:** A549-GFP:Rab11a-FAST:HsMX1-expressing cell infected with A/Victoria/3/75 for 9h30. One image was acquired every 1 second for 30 seconds. GFP:Rab11 is represented in cyan and FAST:HsMX1 in red. The movies are representative of three independent experiments. Scale bar: 10 µm.

**Movie S19.1-19.3:** A549-GFP:Rab11a-FAST:HsMX1-expressing cell infected with A/Victoria/3/75 for 24 h. One image was acquired every 1 second for 30 seconds. GFP:Rab11 is represented in cyan and FAST:HsMX1 in red. The movie are representative of three independent experiments. Scale bar: 10 µm.

**Movie S20.1-20.2:** A549-GFP:Rab11a-FAST:HsMX1-expressing cell infected with A/Victoria/3/75 for 6h30 and imaged every 5 minutes for until 17h15 post-infection. GFP:Rab11 is represented in cyan and FAST:HsMX1 in red. Representation is either an average Z-projection (16.1), or a single slice from the z-stack (16.2). The movies are representative of three independent experiments. Scale bar: 10 µm.
